# Subthalamic Nucleus Optogenetic Inhibition Bidirectionally Regulates Social Motivation According to Familiarity and Social Hierarchy

**DOI:** 10.64898/2026.06.30.735581

**Authors:** Lucie Vignal, Mehdi Bancilhon, Christophe Melon, Yann Pelloux, Nicolas Maurice, Christelle Baunez

## Abstract

**Background:** Social behavior is a core component of mental health, and its disruption characterizes many neuropsychiatric disorders such as autism. Social operant paradigms enable the quantification of volitional aspects of social motivation and interactions. While sex differences have been shown to influence social motivation, factors such as familiarity and social hierarchy are also likely to play a critical role that remain insufficiently explored. In addition, the subthalamic nucleus (STN), traditionally studied in motor circuits, has emerged as an important regulator of reward and motivational processes and may contribute to social behavior processes.

**Methods:** In this study, we examined the influence of peer familiarity (cagemate vs. stranger) and social hierarchy (dominant vs. subordinate) 1) on operant volitional social interaction using a fixed ratio 1 (FR1) and 2) on social motivation using a progressive ratio (PR) schedule of reinforcement in non-isolated rats. To assess the causal contribution of the STN, we used optogenetic photo-inhibition during both tasks in male rats.

**Results:** Male rats displayed a reduction of social interest and motivation toward familiar peers, mainly driven by the social hierarchy, while female did not. STN photo-inhibition in males abolished the familiarity-driven reduction under FR1 but decreased motivation independently of familiarity or hierarchy in PR.

**Conclusions:** These findings highlight sex, familiarity, and hierarchy as key modulators of volitional social behavior and demonstrate a direct role of the STN in regulating social motivation. Together, they provide mechanistic insights into processes that may be disrupted in neuropsychiatric disorders characterized by social dysfunction.

## Introduction

In social species such as humans, social behavior is essential for survival, healthy development, and well-being [1–3]. Disruptions in social reward processing and social motivation are core features of major neuropsychiatric disorders, including autism spectrum disorder (ASD), schizophrenia, and depression. These pathologies often involve diminished initiation of social contact, altered valuation of social stimuli, or reduced effort to obtain social interaction [4,5]. Understanding the mechanisms that govern the motivation to seek social contact is therefore central to elucidating the neurobiology of social dysfunction in psychiatric disorders. New operant paradigms have recently enabled the study of the volitional component of social interactions in rodents. In the social operant conditioning task, rats, a social animal [6], must perform a goal-directed action (e.g., lever pressing) to gain access to a peer through a sliding door and a grid. Mice and hamsters can also acquire social operant conditioning [7–11]. Social interactions reliably act as a reinforcer in this paradigm [12,13], since rodents show greater responding for access to a peer compared to door opening alone, or non-social stimuli [8,14–17]. In rats, several factors modulate social motivation in this paradigm, including housing conditions (isolation vs. pair housing)[18–21], sex of the subject [15,20] and familiarity or sex of the social stimulus [20,22]. Notably, most studies have focused on female rats, who exhibit higher social motivation than males [20]. Moreover females do not show differences in motivation to work for a familiar vs. unfamiliar peer in social motivation paradigm [19,23], while in choice paradigm, they prefer interacting with unfamiliar conspecifics [24]. In contrast, the influence of familiarity and social hierarchy on social motivation in male rats remains largely unexplored, despite their relevance in the studies of social behavior.

Although the operant social motivation paradigm is increasingly used, the neural mechanisms underlying volitional social behavior remain poorly understood. Most work has focused on the reward system, particularly the dopaminergic system [8,9,17,22] as well as the oxytocinergic system [11]. However, converging evidence indicates that the subthalamic nucleus (STN) may also play a key role in social behavior. Indeed, the STN has emerged as a key node in motivational processes, exerting opposite regulation over motivation for palatable food versus drug rewards [25–30]. STN lesions have also been shown to reduce social contacts [31], impair social recognition memory [32], and blunt responses to the emotional valence of ultrasonic vocalizations (USVs), whether positive or negative [33,34]. Similar impairments in emotional experience and recognition, associated with STN activity modulation, have also been reported in humans [35–42]. These findings suggest that the STN contributes both to social interest and to the processing of emotionally salient social cues, without altering responses to objects [32,43]. Furthermore, modulating STN activity alters how social context (peer presence or vocalizations) influences drug use [33,34,43–45]. Thus, dysfunction of STN activity may contribute to the social impairments observed in several neuropsychiatric disorders. Despite this growing evidence, no study has directly tested the causal involvement of the STN in social motivation.

In the present study, we examined how STN optogenetic inhibition modulates operant responding for social interaction and assessed the influence of familiarity (cagemate vs. stranger) and social hierarchy (dominant vs. subordinate) under both low-effort (fixed ratio 1 (FR1)) and high-effort (progressive ratio (PR)) reinforcement schedules.

## Materials and Methods

A detailed description of the methods is provided in the Supplementary Materials. All procedures were conducted in accordance with French regulations (Decree 2010-118) and approved by the local ethics committee and the French Ministry of Agriculture (#44330-2023081011306728 v3). Lister Hooded rats (n = 130; 350 g at arrival; Charles River, France) were either tested behaviorally (n = 68 males, n = 6 females) or used as social peers (n = 56 males). Animals were pair-housed throughout the protocol. Only male rats underwent optogenetic inhibition of the STN. For this, males received stereotaxic injections of either a control virus (AAV5-CaMKII-EYFP; EYFP-control groups) or the inhibitory opsin ARCHT3.0 (AAV5-CaMKII-ArchT3.0-p2A-EYFP; ARCHT3.0 groups) into the STN (coordinates from bregma: AP −3.7 mm, L ±2.4 mm, DV −8.4 mm), with optic fibers implanted 4 mm above the injection site. Females followed the same behavioral procedures as the male cagemate group without surgery or photomodulation. Dominance status was assessed using homecage scoring, the tube test, and a modified food competition test, with rats classified as either dominant (score ≥ 2) or subordinate (score ≤ 1).

Operant behavioral testing took place in customized self-administration chambers with two compartments separated by a sliding door and a grid. One compartment contained two levers with cue lights; pressing the active lever switched on the associated light and opened the door for 30 s, allowing visual, auditory, and partial tactile interaction throught the grid with a peer placed in the second compartment. The peer was either the cagemate or a stranger (one of the eight sex-, age-, and weight-matched unfamiliar male rats rotated daily). Sessions began with fixed ratio 1 (FR1; i.e. continuous reinforcement) training until stable responding (<25% variability across five consecutive sessions). Rats then performed five additional FR1 sessions with laser ON. Photo-inhibition parameters were based on previous electrophysiological and behavioral validation [45]: 532 nm light, 15-s pulses at 0.2 Hz for 5 min, followed by 5 min off, repeated throughout the session. Rats subsequently underwent eight progressive ratio (PR) sessions, with the last four sessions with laser ON, during which lever-press requirements increased progressively (1, 2, 3, 4, 6, 8, 10 …) [18]. Sessions ended after 1h of inactivity or after 4h had elapsed, and the breakpoint was defined as the last completed ratio. The timeline of the behavioral experiment is illustrated in Fig. 1A. Video recordings of some social operant sessions were obtained using an analog tube mini-camera (Active Média Concept) at 24 frames per second (1920×1080 pixels, 1080P), and analyzed using a python script.

**Fig. 1.**
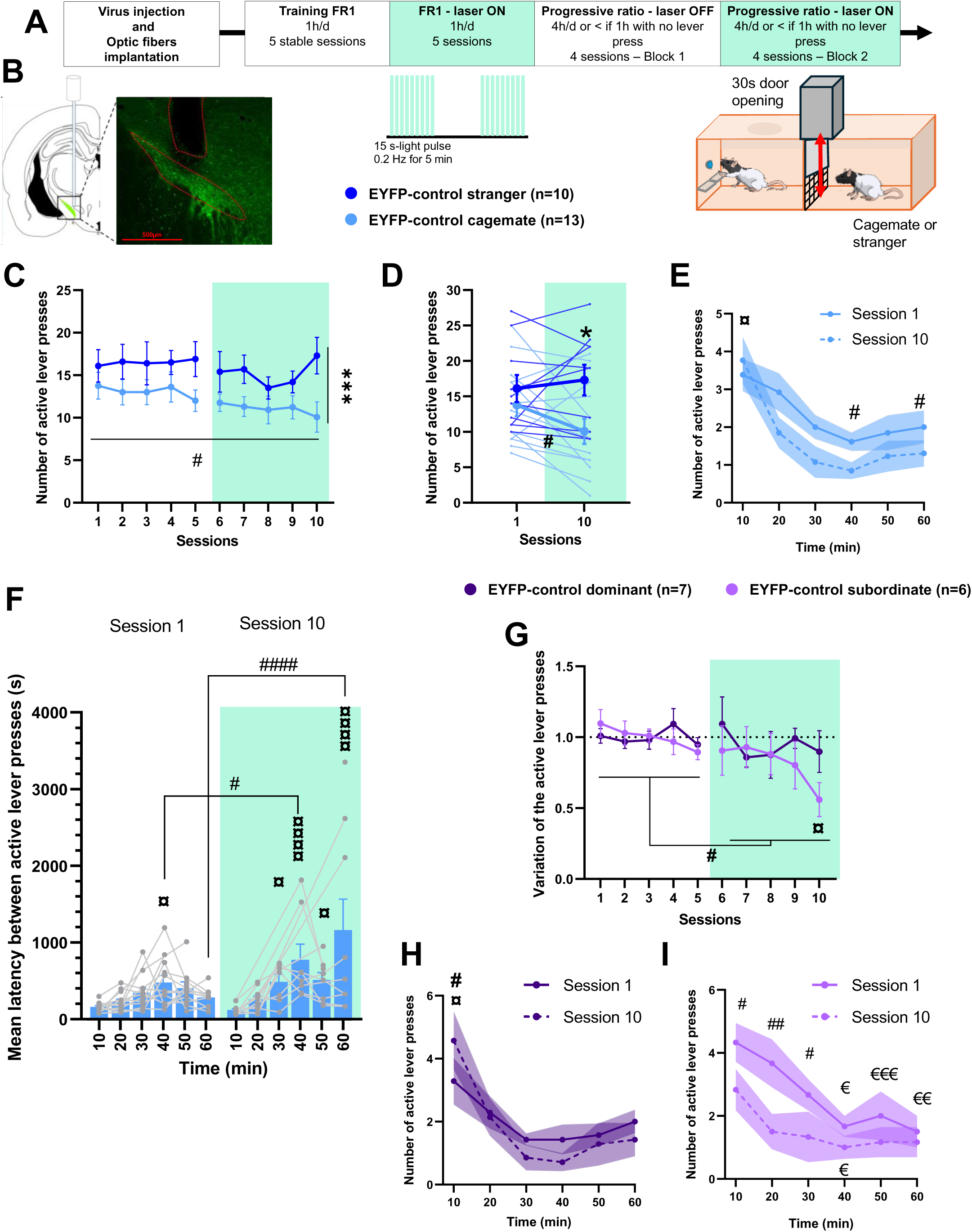
Peer familiarity and social hierarchy in males influence the volitional social interactions in FR1 paradigm. **(A)** Timeline of the behavioral experiment and schematic apparatus. Green boxes represent laser activation **(B)** Schematic coronal section (left) and epifluorescence microscope picture (right) of the STN, after the viral injection and the optic fiber implantation. **(C)** Number of active lever presses (leading to opening the door) per session, performed by the EYFP-control stranger group (interacting with a stranger rat, n=10, dark blue) and by the EYFP-control cagemate group (interacting with their cagemate, n=13, light blue) (peer identity effect (F(1,223)=26.187, p<0.0001, and block effect: F(1,223)=5.451, p=0.0204)). (D) Number of active lever presses during the first and the last session for the EYFP-control stranger group and the EYFP-control cagemate group. **(E)** Number of active lever presses per 10 min-bins within session 1 (plain light blue line) and session 10 (dashed light blue line) in EYFP-control cagemate group (Session effect: (F(1,143)=6.640, p=0.011) and bin effect: (F(1,143)=8.816, p<0.0001). **(F)** Mean latency between each active lever presses per 10 min-bins within session 1 and session 10 (green box) in EYFP-control cagemate group. The grey lines represent the individual data. (bin × session (F(5,102)=4.1032, p=0.0020) (**G)** Variation of active lever pressing normalized with the baseline over the 10 sessions, in EYFP-control subordinate rats (interacting with their dominant cagemate; n=6, pink) and in EYFP-control dominant rats (interacting with their subordinate cagemate; n=7, violet) (block effect: (F(1,107)=8.0337, p=0.0055). (**H)** Number of active lever presses per 10 min-bins within session 1 (plain violet line) and session 10 (dashed violet line) in EYFP-control dominant group (bin × session interaction (F(5,66)=2.7254, p=0.0267). **(I)**. Number of active lever presses per 10 min-bins within session 1 (plain pink line) and session 10 (dashed pink line) in EYFP-control subordinate group (bins (F(5,55)=7.2561, p<0.0001) and sessions (F(1,55)=18.1891, p<0.0001). Pale Green zones : period with laser ON (sessions 6 to 10) * p<0.05, ** p <0.01, *** p<0.001, **** p<0.0001 : between groups (EYFP-control stranger vs EYFP-control cagemate, or EYFP-control subordinate vs EYFP-control dominant) # p<0.05, ## p <0.01, ### p<0.0001 : within sessions or blocks ¤ p<0.05, ¤¤ p <0.01, ¤¤¤ p<0.001, ¤¤¤¤ p<0.0001 : compared to the first bin or session, or first bin being significantly different from the others

Animals with improper viral expression or optic fiber placement were excluded (n = 18), along with one rat whose cagemate died and two rats that failed to learn lever pressing (see **Fig. 1B** for correct expression and implantation). Number of active and inactive lever presses, number of rewards obtained, and latencies between 2 consecutive active lever presses were recorded and analyzed. The number of times the rats went to the door after pressing the active lever, and the corresponding latency, were extracted from video recordings of the last FR1 session and all PR sessions. Statistical analyses were performed using GraphPad Prism 8.0 or R (lmerTest, emmeans, rstatix). Data are expressed as mean ± SEM. Linear mixed-effects models with subject as a random factor were used to analyze lever presses, rewards, and latencies. Two-tailed tests were applied unless specified, with α = 0.05. Results for main figures are presented in **Table 1** and for supplemental figures in **Table 2**.

## Results

All rats kept for the analysis, regardless of group, successfully learned to discriminate active from inactive levers under FR1, reaching performance levels above chance (male: **Fig.S1A-C**, female: **Fig.S2A-C**).

### Volitional social interactions depend on the sex, peer identity and the social hierarchy

#### Sex and peer’s familiarity influence social reward overtime

First, we investigated the influence of familiarity of the peer on the operant responding for access to social interaction in controls rats. For males, the social partner was either a stranger or a cagemate, while in females, only cagemates were tested.

Both groups of EYFP-control male rats associated with a familiar or stranger peer reliably learned to lever press for 30s social interactions, regardless of conspecific identity during baseline sessions. However, male rats pressed more to interact with a stranger than with their cagemate in both blocks of five sessions (**Fig.1C)**. Moreover, while male EYFP-control rats interacting with stranger maintained stable responding over time, those interacting with cagemate presented a progressive decrease of active lever across sessions, with a significant drop from session 1 to 10 (**Fig.1D**).

To better understand how familiarity influences the dynamic of active lever pressing within a session, we divided the sessions 1 and 10 into 10-minute bins and compared them.

In cagemate pairs, in session 1, rats pressed more the active lever during the first 10 minutes than in later bins **(Fig. 1E).** In session 10, pressing was reduced at minutes 40 and 60 compared to session 1, indicating that the overall decrease was driven by late-session disengagement. Analysis of latency between consecutive active lever presses in 10-minute bins confirmed this pattern. Indeed, latency steadily increased after 30 minutes in session 10, reaching significant differences at 40 and 60 minutes compared with the first session. **(Fig. 1F)**. Thus, male rats interacting with cagemate showed progressive disengagement in both frequency and timing of lever pressing, indicating a familiarity-driven decline volitional social interest.

In females, active lever pressing for the cagemate was significantly higher than in males interacting with the same type of peer and unlike male, they did not exhibit decline over time (**Fig.S2D**). However, their strategy changed in session 10: they worked more during the first 10 minutes than during the rest of the session and compared to the first 10 min of session 1, but took more time between lever presses after 40 minutes **(Fig.S2E-F).**

### Social hierarchy within cagemates influences the rewarding effect of social interactions in males only

To assess whether social dominance modulates lever pressing for cagemate interaction, we classified rats within pairs as dominant or subordinate using the tube test, the sucrose competition, and home cage observations (**Fig.S1D** for score repartition). To reduce intra- and inter-group variability due to smaller group sizes (n=7 for dominant and n=6 for subordinate), we normalized the number of active lever presses of the first five sessions (block 1) for each rat.

Both dominant and subordinate rats pressed the lever to interact with their cagemate. Dominant rats kept a stable level of active lever presses over sessions. On the other hand, subordinate rats presented a significant decrease in lever pressing in the second block compared to the first one (**Fig.1G**), with a reduction between the first and last session. Analyzing lever presses and latency across 10-minute bins revealed a different strategy in dominant and subordinates EYFP-control rats. The pressing of dominants remained stable within session 1, while in the session 10, they worked more in the first 10 min, followed by a marked decline in subsequent bins (**Fig1.H)**. This suggests a shift in strategy towards high initial engagement with rapid within-session decline. Latency analysis supported this pattern **(Fig.S1E)**, with stable latencies in session 1 but significantly increased latencies in session 10 during the second half of the session. Thus, although overall performance remained unchanged, the altered within-session dynamics suggest that dominant rats’ initial interest in their subordinate partner may decline with repeated sessions.

Conversely, subordinate control male rats showed reduced active lever press activity after the first half of session 1 compared to the first 10 minutes **(Fig.1I)**. In session 10, activity was already lower during the first half-session compared to session 1, accounting for the overall decrease in total active lever presses. Similarly, latencies increased progressively in session 10 and were significantly higher in session 10 than in 1 for the last 10 minutes (**Fig.S1F**) These results indicate not only reduced activity, but also slower responding over time and across sessions, reflecting an initial reduced interest for interactions with the dominant within the first session, that further declines with repetitions.

In females, social hierarchy had no effect on overall active lever pressing, bin-wise pressing, or latencies (**Table 3**), suggesting that dominance does not influence willingness to interact with the cagemate in female rats.

### STN photo-inhibition suppresses the familiarity and social hierarchy effect

To test the role of the STN in voluntary social interactions, we bilaterally injected the inhibitory opsin ARCHT3.0 into the STN and applied light stimulation during sessions 6–10 (block 2) to transiently inhibit STN neurons [45].

### STN photo-inhibition has no effect on volitional social interaction with a stranger in continuous reinforcement (FR1)

STN photoinhibition in ARCHT3.0 male rats did not affect the willingness to interact with a stranger (no change on active lever pressing **(Fig.2A)**, nor the number of presses or the latency between presses across 10-minute bins). STN photoinhibition had no effect either on the number of times the rats went to the door after pressing the active lever, nor on the corresponding latency for session 10 **(Fig.S1 G *top* and H *left***).

**Fig. 2.**
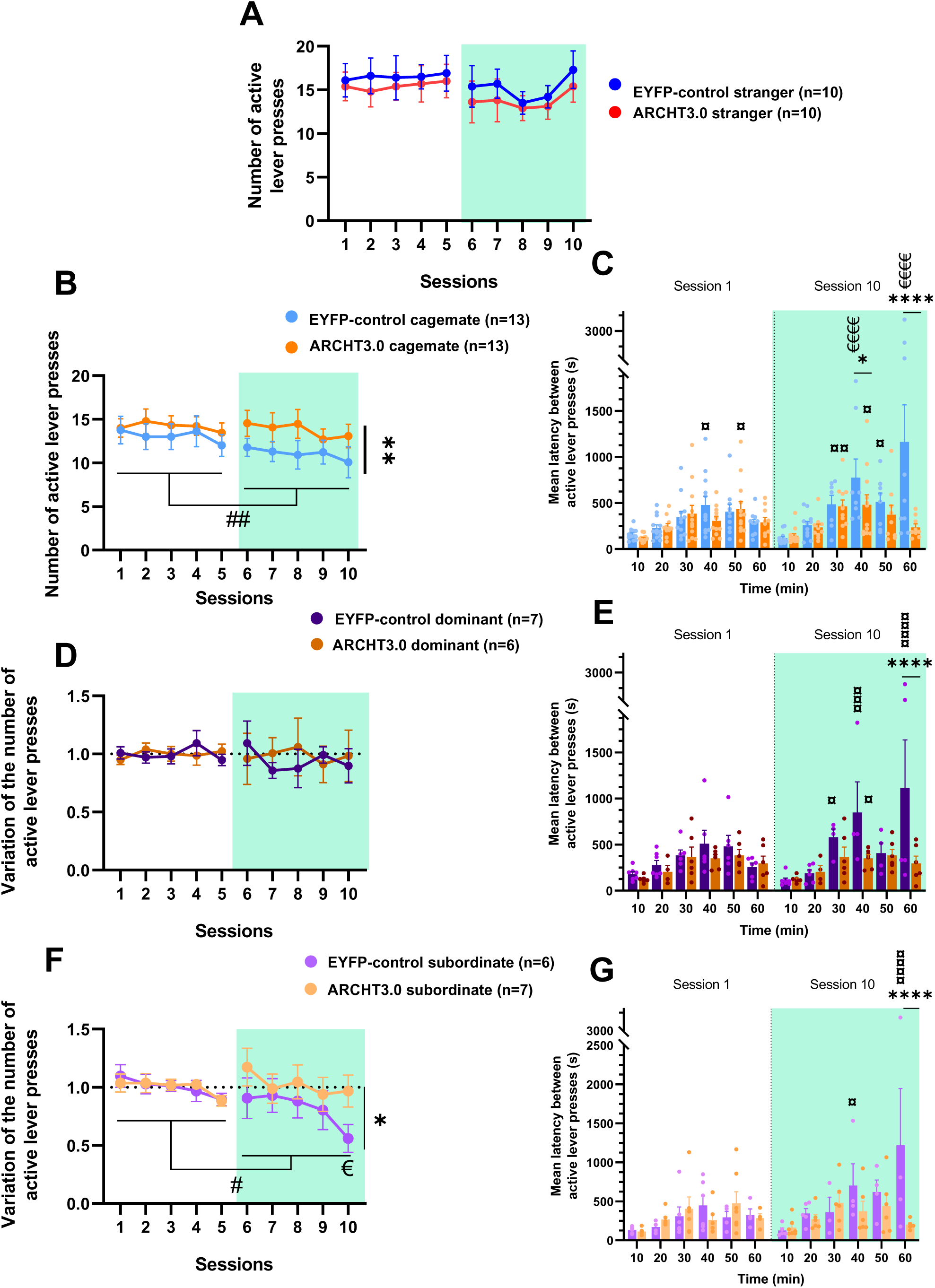
STN photo-inhibition blocks the familiarity- and hierarchy-driven decrease in FR1 schedule. **(A)** Number of active lever presses per sessions, performed to interact with a stranger in control-EYFP (n=10, dark blue) and in ARCHT3.0 rats (n=10, red) (all >0.05) without and with laser ON (green zone). **(B)** Number of active lever presses per sessions, performed by the control-EYFP (n=13, light blue) and the ARCHT3.0 rats (n=13, orange) to interact with their cagemate (Group : F(1,253)=9.568, p=0.0022). **(C)** Mean latency between each active lever presses per 10 min within session 1 (laser OFF, white zone) and session 10 (laser ON (green zone) in cagemate group (Bin x group x session : F(5,213)=3.441, p=0.0052). **(D)** Mean variation of the number of active lever presses normalized by the baseline per sessions, realized by the dominant rats in control-EYFP group (n=7, violet) and in ARCHT3.0 rats (n=6, brown) (all >0.05). **(E)** Mean latency between each active lever presses per 10 min within session 1 (laser OFF, white zone) and session 10 (laser ON (green zone)) in the dominant group (Bin x session: F(5,97)=2.675), p=0.026). **(F)** Mean variation of the number of active lever presses normalized by the baseline per sessions, realized by the subordinate group in control-EYFP (n=6, pink) and in ARCHT3.0 rats (n=7, light orange) (laser x group : F(1,107)=4.648), p=0.0333). **(G)**. Mean latency between each active lever presses per 10 min within session 1 (laser OFF, white zone) and session 10 (laser ON (green zone)) in subordinate group (group x session : F(1,96)=6.433, p=0.0128). Green square : period with laser ON (block 2 : session 6 to 10) * p<0.05, ** p <0.01, *** p<0.001, **** p<0.0001 : between groups (EYFP-control vs ARCHT3.0) # p<0.05, ## p <0.01, ### p<0.0001 : within sessions or blocks ¤ p<0.05, ¤¤ p <0.01, ¤¤¤ p<0.001, ¤¤¤¤ p<0.0001 : compared to the first bin or session

### STN photo-inhibition blocks the familiarity and social hierarchy effects in FR1

In male rats, STN photoinhibition in ARCHT3.0 suppressed the familiarity-driven decrease seen in EYFP-control males interacting with cagemate. This led to a significant group difference during laser activation, but a stable level in the ARCHT3.0 group (block 2 compared to block 1) **(Fig.2B)**. Bin analysis confirmed that ARCHT3.0 rats maintained similar active lever pressing across bins in session 1 and 10 (**Fig.S1I**), unlike EYFP-controls. In ARCHT3.0 rats, latency was transiently increased in bins 3 and 4 compared to bin 1 during session 10, with no significant differences when compared to session 1 (**Fig.2C**). In contrast, EYFP controls displayed a sustained increased latency across bins in session 10, which led to a significant group difference at bins 4 and 6. These results indicate that STN photoinhibition maintained social interest for the cagemate over time and repeated sessions thus preventing the familiarity-driven decline in pressing and associated increased latencies. STN photoinhibition had no effect on the number of times the rats went to the door after pressing the active lever, nor on the corresponding latency for session 10 **(Fig.S1G *bottom* and H *right***).

Next, we examined the effect of STN photo-inhibition on social hierarchy influence in familiarity-driven reduction (score repartition of ARCHT3.0 rats: **Fig.S1J**).

In dominant rats, STN photoinhibition had no effect on overall active lever pressing, with ARCHT3.0 rats remaining stable across blocks and similar to controls (**Fig.2D**). However, latency analysis revealed that STN photoinhibition suppressed the progressive and persistent increases in latency seen in session 10 in EYFP-control rats **(Fig.2E)**. ARCHT3.0 rats only showed a transient rise around the 40th minute (vs. bin 1), resulting in a between-group difference in the last bin of session 10. Thus, although total lever pressing was unaffected, STN photoinhibition altered its dynamics, preventing the familiarity-related slowing observed in dominant controls.

In subordinate rats, STN photoinhibition suppressed the reduction in total active lever presses observed in subordinate EYFP-control rats **(Fig.2F)**. ARCHT3.0 subordinate males maintained a stable number of active lever presses across sessions. This was confirmed by the suppression of the rise of the latencies, resulting in a significant between-group difference in the last 10 minutes **(Fig.2G)**.

Overall, these results indicate that STN photoinhibition abolishes familiarity-driven reductions in voluntary social interactions, irrespective of social hierarchy.

### The motivation to obtain social interactions depends on the sex, the identity of the peer and the social hierarchy

After the FR1 protocol, rats performed a progressive ratio (PR) task to increase the work load required to access either a cagemate or a stranger conspecific, thus further assessing their motivation for social interactions.

In male EYFP-control rats, familiarity influenced the number of active lever presses **(Fig.S3A)** and rewards earned (and therefore the breakpoint). Control males worked less to interact with their cagemate than with a stranger in both blocks **(Fig. 3A)**, earning fewer rewards (stranger: 7.3 (mean of the 4 sessions of block 1) and 6.7 (mean of the 4 sessions of block 2); cagemate: 6.3 and 5.7 rewards in blocks 1 and 2, respectively), corresponding to breakpoints of 11.2 and 9.4 presses for stranger conditions, and 8.6 and 7.4 presses for cagemate conditions. “The cagemate group” also showed a significant decline in lever pressing and rewards obtained across blocks 1 and 2. Dividing each session into four 1-hour bins showed that both groups were most active during the first hour (**Fig.3B**), with consistently higher pressing in the “stranger group”. In block 1, motivation to interact with a stranger showed only a brief dip in hour 3, while that to interact with cagemate showed a progressive decline. In block 2, pressing for interaction with a stranger stabilized after hour 1, but motivation for interaction with cagemate decreased during the first and second hour relative to block 1. Thus, an early between-group difference was amplified by a time-dependent decline in pressing for cagemate rats. Reward counts per 1-hour bin confirmed these findings **(Fig.S3B).** To better capture within-session changes in motivation, we focused on the most active hour (∼5 rewards on average) and compared the latency between the last press of one ratio and the first press of the next, another measure of motivation. This measure reflects how quickly rats re-engage in the task after accessing the social stimulus. Rats exhibited a different pattern depending on the identity of the rat behind the grid/door **(Fig.3C)**: in block 1, latency to re-engage after interaction with the cagemate increased progressively (significant between rewards 1 and 4–5), whereas it remained stable after interaction with a stranger rat. In block 2, this increase appeared earlier (by reward 3) after interaction with cagemate, while there were only minor fluctuations (reward 1 vs. 3 and 5) after interaction with a stranger. Thus, motivation to interact with a stranger rat did not diminish, despite a slight habituation, whereas motivation to interact with the cagemate declined across session blocks.

**Fig. 3.**
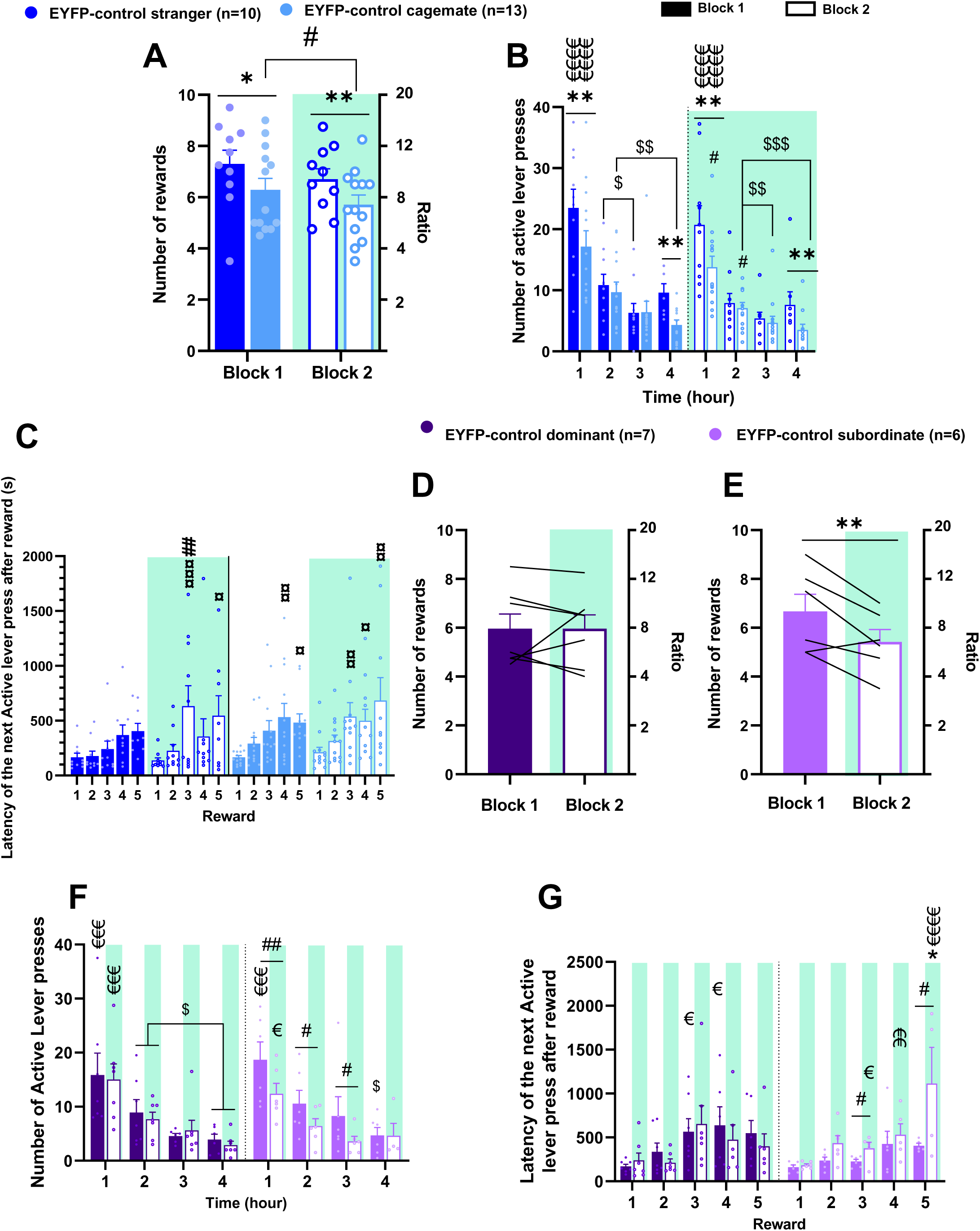
Peer familiarity and social hierarchy in males reduce the social motivation in PR. **(A)** Number of rewards obtained and corresponding ratio completed in 4-session blocks (block 1: laser OFF, plain bar, block 2: laser ON (green zone, empty bar)) in EYFP-control stranger (n=10, dark blue) and EYFP-control cagemate (n=13, light blue) groups (block effect: F(1,177)=4.314, p=0.0392, familiarity effect: F(1,177)=11.855, p=0.0007). **(B)** Number of active lever presses per 1h within block 1 (laser OFF, plain bar) and block 2 (laser ON (green zone), empty bar) (block effect: F(1,649)=11.631, p=0.0007, Bin x familiarity: F(3,649)=6.738, p=0.0002)) **(C)** Latency to make the next active lever press after obtaining a social reward depending on the reward rank (1^st^, second, 3^rd^, etc..)(effect of the reward rank on the latency: F(4,816)=12.640, p<0.0001). **(D)** Number of rewards obtained and corresponding ratio completed in 4-session blocks in dominant EYFP-control group (n=7, violet) (all >0.05). **(E)** Number of rewards obtained and corresponding ratio completed in 4-session blocks in subordinate EYFP-control group (n=6, pink) (block effect: F(1,42)=5.530), p=0.235). **(F)** Number of active lever presses per 1h within block 1 (laser OFF, plain bar) and block 2 (laser ON (green zone), empty bar)) (hierarchy x block : F(1,364)=6.698, p=0.01, bin effect: F(3,364)=61.028, p<0.0001). **(G)** Latency to make the next active lever press after obtaining a social reward in dominant and subordinates (hierarchy x block : F(1,445)=8/242, p=0.0043, rank of the reward: F(4,445)=9.209, p<0.0001). Green square : period with laser ON (block 2: session 4 to 8) * p<0.05, ** p <0.01, *** p<0.001, **** p<0.0001 : between EYFP-control groups (stranger vs cagemate, dominant vs subordinate) # p<0.05, ## p <0.01, ### p<0.0001 : within sessions or blocks ¤ p<0.05, ¤¤ p <0.01, ¤¤¤ p<0.001, ¤¤¤¤ p<0.0001 : compared to the first bin or session, or first bin being significantly different from the others $ p<0.05: between bins of the same block

To examine the influence of the social hierarchy, EYFP-control “cagemate” rats were split into dominant and subordinate groups. Dominant rats showed no change in motivation across blocks (number of active lever presses (**Fig. S3C)** and number of rewards earned in both block: 6.0 reward, ratio : 8)(**Fig.3D**) to interact with their subordinate. In contrast, subordinate rats significantly decreased their motivation (number of active lever presses (**Fig.S3D**) and number of rewards earned from 6.7 (ratio:9.4) to 5.4 (ratio: 6.8) (**Fig.3E**)), to interact with their dominant from block 1 to block 2 (**Fig.3G**). When comparing the number of lever press per hour, both groups were most active in the first hour (**Fig. 3F**), with lever pressing declining over time in block 1. In block 2, dominant rats maintained their block 1 pattern of responses, whereas subordinates pressed less during the first three hours compared to block 1.

Dominant rats were slower to press the active lever after rewards 3 and 4 in block 1 (vs. reward 1), but showed stable latencies in block 2. Subordinate rats displayed stable latencies in block 1 but slowed their responding in block 2 from reward 3 onward compared to reward 1, to block 1, and to dominants after reward 5 (**Fig.3G**). Overall, unlike dominants, subordinates showed a time-dependent decline in engagement to interact with their dominant.

In females, the motivation to interact with the cagemate remained constant between blocks (no significant changes in rewards earned: 8.6 (ratio: 14.4) vs. 8.2 (ratio: 12.8) **(Fig.S2G)**, active presses **(Fig.S2H)**). In both blocks, females worked more during the first hour **Fig.S2I-J**) but remained at a steady stable level of pressing between blocks, unlike males. There was no influence of the hierarchy (dominant vs. subordinate) in females on the motivation, as assessed by the number of rewards obtained (dominant: 7.1 (ratio: 10.2) and 7.25 rewards (ratio:10.5); subordinate: 10.1 (ratio: 14.2) and 9.1 rewards (ratio: 16.2).

### STN photo-inhibition reduces social motivation irrespective of peer type in PR

During sessions 5 to 8 (block 2), the laser was turned on to induce STN photo-inhibition in ARCHT3.0 rats. In ARCHT3.0 male rats working to interact with a stranger, motivation was comparable to controls when the laser was off (block 1). However, STN photo-inhibition (laser ON, block 2) reduced the motivation (i.e. number of active lever presses **(Fig.S3E)** and the number of rewards earned (and therefore the breakpoint: from 6.9 (ratio:9.8), to 5.7 (ratio:7.4) **(Fig.4A))**. Indeed, STN inhibition decreased pressing during the first hour compared to controls and to the laser-off condition, and during the third and last hours compared to laser-off **(Fig.4B)**, suggesting a sustained reduction in motivation. To test whether STN inhibition altered response timing, we compared the latency to perform the first press of the session (before identification of the partner), the second (after identification of the partner), and last lever press (before the rat stopped working, reflecting persistence in responding). In ARCHT3.0 rats, the last press of the session occurred earlier during STN photo-inhibition **(Fig.4C)**, indicating that they stopped working earlier than in laser-off conditions. Overall, STN inhibition decreased willingness to work for social interaction with a stranger. However, STN photo-inhibition did not modify the number of times the rat went to the door following its opening, nor the latency to approach. (**Fig.S3H *top* -I *left***).

**Fig. 4.**
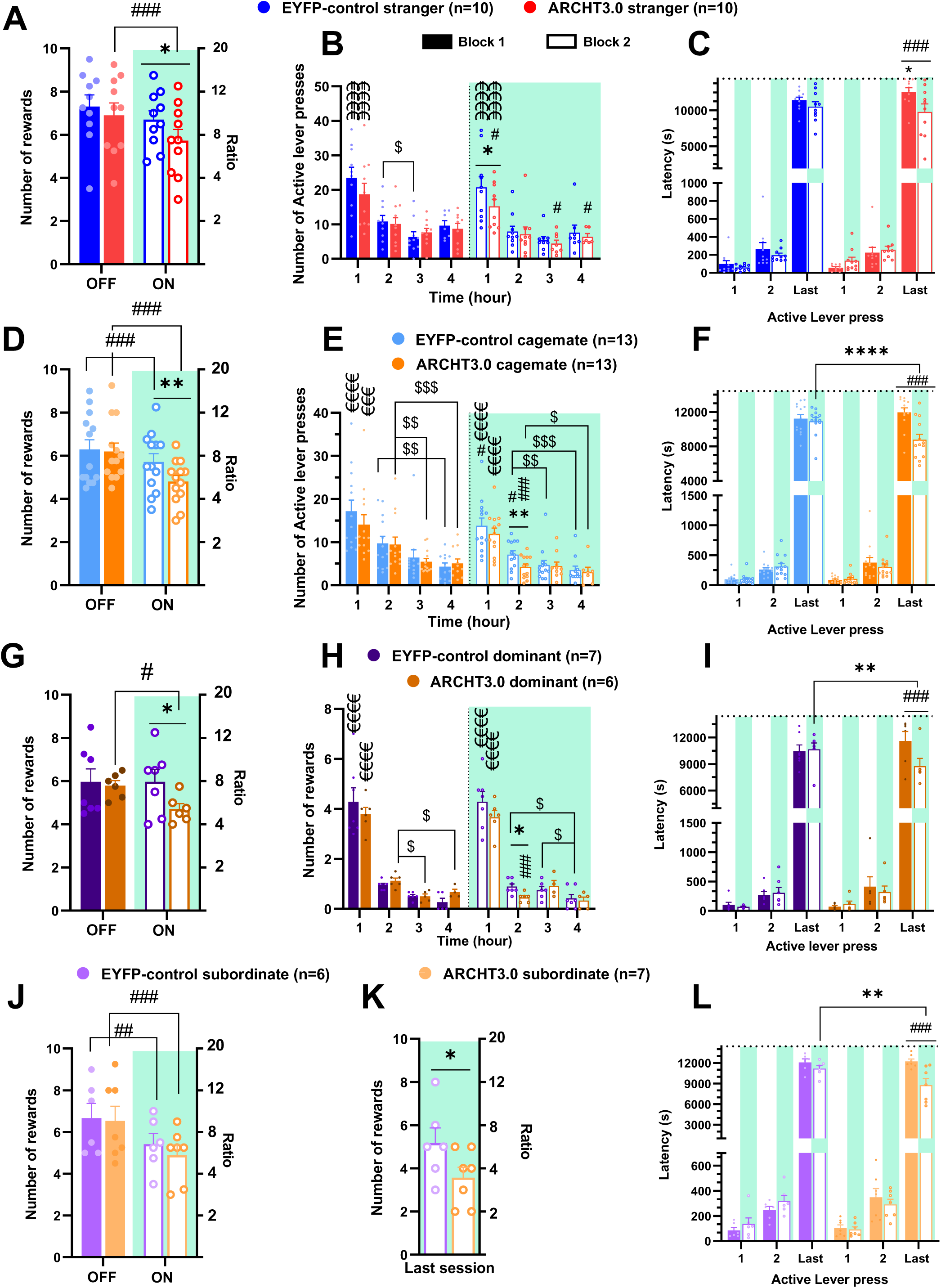
STN photo-inhibition decreases social motivation in PR task irrespective of the familiarity and social hierarchy. **(A)** Number of rewards obtained and last ratio reached in 4-session blocks (block 1: laser OFF, plain bar; block 2: laser ON, empty bar, green zone) in EYFP-control stranger (n=10, dark blue) and ARCHT3.0 stranger (n=10, red) groups (laser: F(1,153)=8.306, p=0.0045). **(B)** Number of active lever presses per 1h within block 1 and block 2 in strangers groups (bin x group : F(3,552)=3.181, p=0.0237, laser effect: F(1,552)=12.945, p=0.0003). **(C)** Latency to do the first, second and last lever press in “stranger” groups (laser x group x active lever press: F(2,447)=3.037, p=0.0490). **(D)** Number of rewards obtained and corresponding last ratio completed in 4-session blocks in EYFP-control cagemate (n=13, light blue) and ARCHT3.0 cagemate (n=13, orange) groups (laser effect: F(1,201)=15.891, p<0.0001, group effect: F(1,201)=4.130, p=0.0434). **(E)** Number of active lever presses per 1h within block 1 and block 2 in “cagemate” groups (bin effect : F(3,748)=81.022, p<0.0001, group effect: (F(1,748)=6.570, p=0.0106) **(F)** Latency to do the first, second and last lever press in block 1 and in block 2 in cagemate groups (laser x group x active lever press: F(2,587)=7.986, p=0.0004). **(G)** Number of rewards obtained and corresponding last ratio completed in 4-session blocks in EYFP-control dominant (n=7, violet) and ARCHT3.0 dominant (n=6, brown) groups (group: F(1,97)=4.239, p=0.0422). **(H)** Number of social rewards obtained per 1h within block 1 and block 2 in dominant groups (bin x group : F(3,327)=3.076, p=0.0278). **(I)** Latency to do the first, second and last lever press in block 1 and in block 2 in dominant groups (laser x group x active lever press: F(2,288)=3.960, p=0.0201). **(J)** Number of social rewards obtained and corresponding last ratio completed in 4 sessions blocks in EYFP-control subordinate (n=6, pink) and ARCHT3.0 subordinate (n=7, light orange) groups (laser: F(1,201)=15.891, p<0.0001, group : F(1,201)=4.130, p=0.0434). **(K)** Number of social rewards obtained and corresponding last ratio completed during the last session of PR in subordinate groups (one tailed p=0.0406). **(L)** Latency to do the first, second and last lever press in block 1 and in block 2 in subordinate groups (laser x group x active lever press: F(2,299)=3.606, p=0.0283). Green square : period with laser ON (block 2 : session 5 to 8) * p<0.05, ** p <0.01, *** p<0.001, **** p<0.0001 : between groups (EYFP-control vs ARCHT3.0) # p<0.05, ## p <0.01, ### p<0.0001 : within sessions or blocks ¤ p<0.05, ¤¤ p <0.01, ¤¤¤ p<0.001, ¤¤¤¤ p<0.0001 : compared to the first bin or session, or first bin being significantly different from the others $ p<0.05 : between bins of the same block

In male rat working to interact with their cagemate, control and ARTCH3.0 group had a similar motivation during the off period. Motivation was reduced during the ON period (lower number of active lever presses (**Fig. S3F**) and rewards earned), with a stronger decrease in ARCHT3.0 STN photoinhibited rats (from 6.1 rewards (ratio:8.2), to 4.8 rewards (ratio:5.6), yielding a significant between-group difference **(Fig.4D)**. Hour-by-hour analysis revealed that ARCHT3.0 rats pressed significantly less during the second hour of laser ON period compared to controls and to their own OFF period **(Fig.4E)**. ARCHT3.0 rats ceased responding earlier than controls (shortened latency to the last active press of the session) **(Fig.4F)**. These results indicate that STN photo-inhibition also reduces social motivation for the cagemate. STN photo-inhibition did not modify the number of times the rat went to the door after its opening, nor the latency to approach. (**Fig.S3H *bottom* -I *right***).

Since STN optoinhibition reduced motivation to interact with the cagemate, the influence of the hierarchy was then analyzed. In dominant rats, STN-photoinhibition decreased the motivation to interact with their subordinate (reduced number of rewards obtained in ARCHT3.0 groups from 5.7 (ratio:7.4), to 4.7 (ratio:5.4)) **(Fig.4G)**. This reduction was particularly marked during the second hour of block 2 compared to the off period and the control group **(Fig.4H)**. ARCHT3.0 dominants also stopped working earlier than controls and their own OFF period **(Fig.4I)**.

In subordinate rats, both control and ARCHT 3.0 groups had a decreased motivation to interact with their dominant cagemate in the second block (laser ON) (no group difference) **(Fig.4J)**. However, the STN inhibition enhanced this reduction in the final session of block 2, since ARCHT3.0 rats earned fewer rewards than controls (3.6 vs. 5.2 rewards and 3.6 breakpoint vs 6.4 respectively) (**Fig.4K**). ARCHT3.0 subordinate rats stopped pressing earlier than both controls and their OFF period across block 2 **(Fig.4L)** and in the last session **(Fig.S3G)**. STN photoinhibition thus reduced motivation of subordinate to work to interact with their dominant.

## Discussion

In this study, we characterized volitional social interaction in male and female rats using both continuous (FR1) and progressive ratio (PR) schedules of reinforcement and examined the causal contribution of the STN. We found that familiarity exert a sex-dependent influence: male rats showed reduced willingness to work for access to their cagemate compared with a stranger, along with a progressive decline in responding across time. This effect was absent in females, who displayed overall higher responding toward their cagemate. Further analysis indicated that the familiarity-driven decline in males was primarily attributable to the fact that subordinate rats are less motivated to interact with their dominant cagemate. Dominant rats, by contrast, maintained relatively stable responding, although showing a time-dependent increase in latencies. STN photo-inhibition abolished the familiarity-driven effect in the FR1 sessions for both subordinate and dominant male rats. However, under the PR schedule, when the effort to gain access to the social interaction increases, STN photo-inhibition induced a generalized reduction in motivation, irrespective of familiarity or hierarchy.

### Effect of sex, familiarity and social hierarchy on volitional social interactions and social motivation in rats

In male, higher motivation for interaction with a stranger may reflect the greater rewarding value of interacting with a novel peer compared to a familiar one. Indeed, many studies have reported that males prefer to interact with, or to receive social cues from, a novel rather than a familiar peer [34,46–51]. In the present study, although the stranger was not entirely novel (rotations from a pool of eight unfamiliar rats rather than a new individual per session), we observed no reduction in lever pressing across sessions, suggesting that, unlike the habituation seen in cagemates over time, the various partners used remained effectively non-familiar.

Female cagemates showed a higher level of lever pressing to interact with their cagemates than males in the same condition. In operant social conditioning paradigms, some studies report no difference between sex [13,14,19,52–55], whereas others do [15,16,20,56]. Most of these studies involve socially isolated rat, a factor known to increase the rewarding properties of social interaction [18,19,57], which could mask potential sex differences. Females also appear less sensitive to the cost associated with social interaction [20] and, in non-isolated conditions, show no difference in motivation toward familiar versus stranger conspecifics [19,23], yet they prefer strangers over familiar when given a choice [24]. Under non-isolated condition, females may be more motivated to interact overall, while the preference for a stranger peer may be stronger in males [49]. Males may also be more sensitive to the identity of the partner, showing higher motivation to interact with a female than with a male, while females’ motivation is unaffected by the partner’s sex [22].

In males, we also observed a hierarchy-dependant effect within the cagemates, driven by subordinates. Such effect was not reported in females, however, future studies with larger sample sizes are needed to confirm this. The hierarchy-driven effect in males may result from a higher rewarding value that dominants attribute to interacting with their subordinate, than subordinate toward their dominant. Indeed, interactions with the cagemate induce CPP only in dominant rats, not in subordinates [43]. In a cocaine self-administration paradigm, the presence of their subordinate significantly reduces drug intake in dominant rats, whereas the presence of a dominant does not alter intake in subordinates, suggesting that the presence of the subordinate acts as an alternative reward to cocaine [43]. This suggests that dominance hierarchy may modulate the quality of social interactions. Indeed, in a prosocial choice task, in which one rat can choose to deliver a palatable pellet to its cagemate without gaining self-benefit for itself, subordinates displayed more social cues than dominants, However, dominants seemed more responsive to these cues, spending more time sniffing through the wall than subordinates. As a result, they acquired the task more quickly and made prosocial choices more frequently than subordinates [58]. Additionally, dominants spend more time investigating unfamiliar rats than subordinates [59], suggesting greater social interest toward both familiar (if subordinate) and unfamiliar partners. A mutual operant social interaction task, where both rats must press a lever to interact [60], could provide evidence that subordinates are less willing than dominants to work for social contact. Thus, as we showed, accounting for social hierarchy is essential in studies of male social interactions, as subordinates and dominants display distinct motivational profiles.

### STN-photoinhibition has opposite effect on volitional social interaction in continuous reinforcement and in progressive ratio schedule

In the FR1 paradigm, where effort requirements are minimal, STN photo-inhibition prevented the decrease in lever pressing in subordinate rats and the increase in inter-press latency in both subordinate and dominant rats. This likely reflects a disruption of cagemate recognition or habituation. Indeed, STN lesions reduce certain type of social contacts [31], and the remaining interactions may be insufficient to support peer recognition. In addition, in our grid-based set-up, access to physical and olfactory cues (anogenital contacts) are limited [48]. Consistently, STN inactivation prevents habituation to repeated peers, abolish recognition of a previously encountered rat [32] and impairs discrimination of USVs from strangers or cagemates [34]. It also allows to develop CPP for a compartment paired with a dominant peer, an effect absent in controls [43]. Thus, in our task, STN inactivation prevents the reduction of interest normally induced by familiarity, by blocking the social habituation or recognition. This effect could also explain the suppression of peer identity-related modulation of cocaine intake [43].

An alternative explanation would be an increase in the hedonic value of social interaction *per se*, without affecting social memory. However, we did not observe an increase in active lever pressing relative to baseline in photo-inhibited rats, in either the stranger or cagemate condition, nor any changes in the latency to approach the door once it opened. This suggests that STN inactivation primarily affected motivational drive rather than the consummatory phase of social interaction. In others studies, STN inactivation had no effect on FR1 responding for food, cocaine, or alcohol [25,26,29,27,30]. Thus, it is more likely that STN photoinhibition prevented social habituation or recognition than the hedonic value of social interaction.

In the PR task, STN photo-inhibition induced a general decrease in motivation, independent of familiarity or hierarchy. Although this contrasts with FR1 results, previous studies also report divergent effects of STN modulation on FR1 and PR schedules of reinforcement. While there is no effect in FR1, modulating STN by lesion or deep brain stimulation in PR task reduced motivation for cocaine and increased it for food [25–27]. This likely reflects the different processes engaged: PR schedules require cost–benefit evaluation, in which the STN is involved, whereas FR1 does not [61–63]. Indeed, STN neuronal subpopulations encode both the required effort and the expected reward, and then integrate them into a subjective motivational value [64,65]. Their relative activity could guide cost–benefit decisions [64,65]. Photoinhibition could disrupt this balance, preventing proper evaluation of effort versus benefit.

The reduced motivation for social interaction may also explain why STN inactivation overrides the protective effect of social context (peer presence or USV playback) on drug intake [33,34,44,45]. With lower social interest, rats are less sensitive to social context and maintain the same level of drug use.

Thus, regardless of the required effort, STN inactivation modifies the volitional aspect of social interaction, highlighting a critical role for the STN in social behavior. In humans, the STN has also been linked to facial and emotional decoding of others or the self [40,42,66,67], and STN deep brain stimulation in parkinsonian patients can be associated with impairments in social behavior [68,69]. Dysfunction of the STN could contribute to social deficits observed in neuropsychiatric disorders, including autism spectrum disorder, schizophrenia, and depression, as well as to potential social side effects of deep brain stimulation in Parkinson’s disease. Understanding how the STN integrates social information with motivational processes could provide key insights into the neural mechanisms underlying social dysregulation and identify potential therapeutic targets. Future studies should further dissect its role within the broader social brain circuitry.

## Acknowledgments

We thank the institute’s platforms involved (PAG, NIT, S-prim, MPRC), as well as Flora Vinet and José Michele Dören for their assistance with the behavioral experiments. AI-assisted technology was used for spelling, typos correction and English polishing.

## Funding

Centre National de la Recherche Scientifique (CB, NM, CM). Aix-Marseille Université (CB, NM)

NeuroMarseille of Aix-Marseille Université (NM)

Institut de Recherche en Santé Public (IRESP-19-ADDICTIONS-02) (CB) Institut National de lutte contre le Cancer (INCA_16032) (LV)

Fédération pour la Recherche sur le Cerveau/ France Parkinson (CB)

## Author contribution

Conceptualization: LV, CB

System development: MB, YP, LV

Performance: LV, CM

Data analyses: LV, MB

Funding acquisition: LV, CB, NM

Writing – original draft: LV, CB, NM

Writing – review & editing: LV, CB, NM

## Data and materials availability

All data are available in the main text or the supplementary materials, or on request to the authors.

