## Supplemental material for "Subthalamic Nucleus Optogenetic Inhibition Bidirectionally Regulates Social Motivation According to Familiarity and Social Hierarchy"

**Supplementary material**

**Materials and Methods**

**Animals**

Lister Hooded rats (n=130; 350 g upon arrival) (Charles River Laboratories, Saint-Germain-sur l'Arbresle, France) were subjected either to behavioral testing (n=68 males, n=6 females) or to be used as peer only (dedicated otherwise to other experiments; n= 56 males) Animals were pair-housed, handled daily and provided *ad libitum* access to water and food (Scientific Animal Food and Engineering, Augy, France), in a temperature-controlled room, under a 12hours inverted light/dark cycle. Experiments were conducted during the dark phase of the cycle (7am-7pm). All procedures complied with the recommendation for animal experiments issued by the European 389 Commission directives 219/1990, 220/1990, and 2010/63, and approved by the local Ethic Committee and the French Ministry of Agriculture (APAFIS #44330-2023081011306728 v3).

**Fiber optic implant design**

Fiber optic implants were built using 230 µm optic fibers (NA 0.22, Thorlabs) glued to 2.5 mm ceramic ferrules (Thorlabs) using epoxy. Ferules were slightly grinded with a dremel to improve the contact with the dental cement.

**Virus**

To transfect STN neurons, we used an AAV5 containing the recombinant protein expression CamKIIa as a promotor control (UNC Vector Core, Chapel Hill, USA). The ARCHT3.0 group received the construct containing the inhibitory opsin AAV5-CaMKII-ARCHT3.0-p2A-EYFP-WPRE, while the EYFP-control animals received it without the opsin, AAV5-CaMKII-EYFP .

**Surgery**

Males rats (~400g) were anesthetized with isoflurane (5%/LO2 for induction then 2-3%/LO2 for 403 maintenance) or a mixture of ketamine (Imalgen, Merial, 100mg/kg, i.p.) and medetominine (Domitor, 404 Janssen, 0.5mg/kg, i.p.) reversed by atipamezole (Antisedan, Janssen, 0.15 mg/kg, i.m.) at the end of the surgery. They were mounted on a Kopf stereotaxic apparatus (Koft instrument) for bilateral STN viral injection and optic fibers implants just above the STN. They received amoxicillin (Duphamox, LA, Pfizer, 100 mg/kg, s.c.), lidocaine (Lurocaïne, Vetoquinol, 20mg/ml, sc and meloxicam (Metacam, Boehringer Ingelheim, 1 mg/kg, s.c.) for antibiotic treatment and analgesia. 0.45 µL of virus were injected bilaterally at 166 nL/mn into STN (in mm: -3.7 AP, ±2.4 L from bregma, -8.40 DV from skull surface, with the incisor bar at -3.3 mm [1]), averaging bregma and interaural coordinates, and optic fiber were implanted at the same coordinate, 4 mm above each injection site. Four screws were anchored into the skull and the system was secured within a dental cement head-cap. Rats were then given a 10 days recovery period.

**Behavioral experiment**

1- **Apparatus**

Experiments took place in homemade self-administration chambers (60x30x31cm), with two compartments, separated by a metallic grid and a homemade automatic guillotine sliding door built with re-used computer CD/DVD player motor and mechanism. One compartment (30x30x31cm) was equipped with two levers, each with a distinct cue light above. They were positioned on the opposite wall to the grid and door. Opening of the door led access to square of grid (13,5x11,5cm) connected with the second smaller compartment (20x30x31cm) in which the social peer could be placed. A custom-built interface and software controlled the operant system (developed by Y. Pelloux, M. Bancilhon and L. Vignal). Video of some social operant sessions were recorded using analogic tube mini-camera (Active Média Concept) at 24 frames per second (1920×1080 pixels, 1080P).

**2- Optogenetic manipulations during behavioral testing**

Only male underwent optogenetic inhibition of STN. Female followed the same behavioral protocol than male working to interact with their cagemate without optogenetic surgery and modulation. Male rats implants were connected to a 200 mW 532 or 528 nm DPSS laser through an optic coupler (FCMM625-50A, Thorlabs) connected to a rotary joint (FRJ_1x1_FC, Doric lenses). Light pulses were controlled by a signal generator (3800 multistim, AM System), with the parameters : 15s light pulse at 0.2 Hz pulse train. Light pulses were discontinuous, alternating 5-min intervals of light ON (at 0.2 Hz) with 5-min intervals of light OFF. Before experiments, light power at the fiber tip was set to 5Mw using a power meter (PM20A, Thorlabs). The photo-inhibition of STN neurons with the opsin ARCHT3.0 with this light pattern was confirmed *in vitro* and *in vivo* in a previous study [2].

**3- Experimental procedure of social self-administration**

The tested rats were placed in the self-administration compartment and the peer (half with their cagemate and the other half with a stranger) in the other compartment. Rats were not isolated before or after the session. Stranger consisted in a daily rotation of 8 sex, weight- and aged-matched peers that they had not encounter before.

Rats were first trained in 1h sessions under a fixed ratio 1 (FR1) schedule of reinforcement, for which one press on the active lever triggered the opening of the door for 30 seconds [3] allowing visual, auditory communication and some physical contacts through the grid with the peer. The cue light was switched on for 20 seconds. Then the door closed automatically and could not be re-opened for 10 seconds (time-out). Any active lever press during the 30s of open door and the 10 seconds time-out had no consequences and was counted as a perseverative response. Lever presses on the inactive lever had no consequences either and were counted as inactive lever presses. The optic fiber were connected to the optic coupler with the laser OFF for cable habituation until stabilized number of active lever press (<25% variability for 5 consecutive sessions) – i.e. baseline. Then rats underwent 5 more sessions, with laser switched ON.

In the second part of the experiment, rats underwent 8 sessions of progressive ratio (PR) schedule of reinforcement, in witch the requirement on the active lever press to open the door for 30s increased progressively according to the following series: 1, 2, 3, 4, 6, 8, 10, 12, 16, 20, 24, 28, 32 … [4]. The self-administration session terminated when the rat stopped pressing the lever for 1h or after 4h had elapsed. The breakpoint was defined as the last completed ratio. Male rats did 4 sessions of PR ratio without laser and 4 sessions with the laser ON.

**4- Evaluation of hierarchy between pairs of rats**

The dominance status within each homecage (i.e. cagemate condition) was assessed by making the addition of score of the three following tests:

- behavioral scoring in the homecage (number of pinning, pouncing, push etc..) during 10 min. The dominant rat scored 1, while subordinate scored then 0.
- the tube test [5,6]. One rat was placed at each extremity of the transparent Plexiglas tube. The diameter of the tube does not allow rats to pass through simultaneously. One

has to reverse its direction. The one retreating was designated as the loser and considered the subordinate (score 0). The test was replicated three times for each pair.

- The modified food Competition [7] : this test took place in a Perpex box (90 x 35 x33 cm), with 9 pellets of sucrose put in a cup in the middle of the box. Only one rat could have access to the cup. We video-recorded the session to *a posteriori* determine which rat ate the most (always the same), and was designated as the dominant (score 1).

Rats were classified as dominant if their score was ≥2, and as subordinate if their score was ≤1.

**Histology**

At the end of experiments, rats were deeply anesthetized (ketamine (200 mg/kg) and medetomidine (60 mg/kg, ip) and then perfused intra-cardiacally with a 4% paraformaldehyde in PBS. Brains were extracted, cryo-protected in sucrose (30%), frozen and sectioned in coronal slices (40 µm thickness) with a cryostat. Then brain section were mounted on glass slides with immunomont (Epredia) and examined with an epifluorescence microscope (Zeiss, Imager.z2). Animals with improper fluorescence localization or optic fiber misplacement were excluded from the results (n=18 in total). Representative correct optic fibers' placement and fluorescence expression are illustrated in Fig. 1A. 1 rat was excluded because his cagemate died, and 2 others because they never learned to lever press.

**Data processing**

The number of rewards obtained, active and inactive lever presses, and their timing of occurrence were extracted from our data and analyzed. Latencies between lever presses were calculated such as the difference between two consecutive active lever presses. The discrimination ratio was calculated as the number of presses on the active lever (sum of active and perseverative presses) divided by the total number of lever presses (sum of active, perseverative, and inactive presses). The number of times the rats went to the door after pressing the active lever, and the corresponding latency, were extracted from video recordings of the last FR1 session and all PR sessions, and annotated using a custom Python interface.

**Statistical analysis**

All variables are expressed as mean ± SEM, statistical tests were two-tailed (unless specified), with a significance threshold set at $\alpha$=0.05. Statistical tests and graphs were performed using GraphPad Prism 8.0 or R (package lmerTest , emmeans and rstatix ). The number of rewards, active or inactive lever presses, and latencies were analyzed by using a linear mixed-effects model with a random intercept. Optogenetic groups (EYFP-control or ARCHT3.0) and condition (peer's familiarity: cagemate or stranger) were considered as between factors, and sessions, or period (laser OFF (block 1) or ON (block 2)), bin (10 min (1 to 6) or 1h-bin (1 to 4)) or active lever press (first, second or last) as within factors, and the subjects as random factor. Analyses were followed by post-hoc analyses. Analysis of results are presented in table 1 for the main figures and in table 2 for supplemental materials.

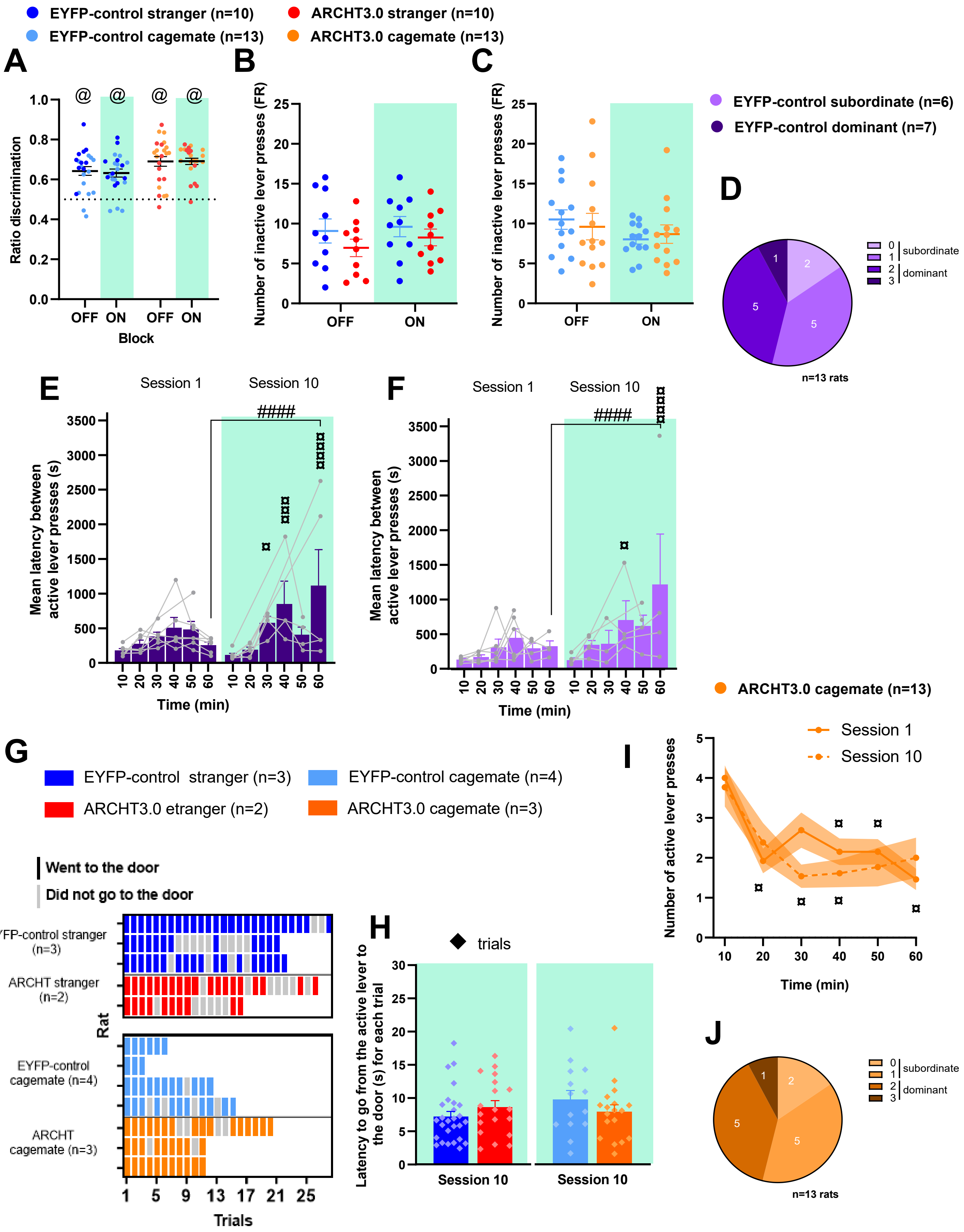

EYFP-control stranger (n=10)
ARCHT3.0 stranger (n=10)
EYFP-control cagemate (n=13)
ARCHT3.0 cagemate (n=13)
A
Ratio discrimination
OFF ON OFF ON
Block
B
Number of inactive lever presses (FR)
OFF ON
C
Number of inactive lever presses (FR)
OFF ON
EYFP-control subordinate (n=6)
EYFP-control dominant (n=7)
D
subordinate
dominant
n=13 rats
E
Session 1
Session 10
Mean latency between active lever presses (s)
Time (min)
F
Session 1
Session 10
Mean latency between active lever presses (s)
Time (min)
ARCHT3.0 cagemate (n=13)
G
EYFP-control stranger (n=3)
EYFP-control cagemate (n=4)
ARCHT3.0 etranger (n=2)
ARCHT3.0 cagemate (n=3)
Went to the door
Did not go to the door
EYFP-control stranger (n=3)
ARCHT stranger (n=2)
EYFP-control cagemate (n=4)
ARCHT cagemate (n=3)
Rat
Trials
H
trials
Latency to go from the active lever to the door (s) for each trial
Session 10
Session 10
I
Session 1
Session 10
Number of active lever presses
Time (min)
J
subordinate
dominant
n=13 rats

**Fig. S1: STN role on familiarity and social hierarchy in males in FR1**

**(A)** Discrimination ratio (active lever presses (active + perseverative) / by total lever presses (active + perseverative + inactive) in FR1 in all groups (EYFP-control stranger n=10, dark blue, cagemate n=13, light blue, ARCHT3.0 stranger : n=10, red, cagemate: n=13, orange) **(B)** Number of inactive lever presses in FR1 in rats working to interact with stanger (all >0.05) **(C)** Number of inactive lever presses in FR1 in rat working to interact with cagemate (all >0.05) **(D)** Distribution of dominance score in EYFP-control cagemate rats **(E)** Mean latency between each active lever presses per 10 min within session 1 (laser OFF, white zone) and session 10 (laser ON (green zone)) in EYFP-control dominant group (Bin x session: F(5,50)=2.932), p=0.0212). **(F)** Mean latency between each active lever presses per 10 min within session 1 (laser OFF, white zone) and session 10 (laser ON (green zone)) in EYFP-control subordinate group (session: F(1,40)=8.728), p=0.0052). **(G)** Individual number of trials in which rat went to the door after an active lever press (colored bar) or did not (grey bar) during session 10 in rats working to interact with stranger (top, group: ($\chi^2(1)$= 0.89, p=0.35) or with cagemate (bottom, group: group: ($\chi^2(1)$<0.0001, p=1.00). **(H)** Mean latency for rats to go from the active lever to the door after it opened (each trial represented by a diamond), for groups interacting with stranger (left, all >0.05) or with cagemate (right, all>0.05) **(I)** Number of active lever presses per 10 min-bins within session 1 (plain orange line) and session 10 (dashed orange line) in ARCHT3.0 cagemate group (Bin: F(1,143)=8.288, p<0.0001) **(J)**. Distribution of dominance score in ARCHT3.0 cagemate rats

@ p<0.0001: significantly different from hazard (50%)
* p<0.05, ** p <0.01, *** p<0.001, **** p<0.0001 : between groups (EYFP-control vs ARCHT3.0)

### p<0.05, ## p <0.01, ### p<0.0001 : within sessions or blocks

¤ p<0.05, ¤¤ p <0.01, ¤¤¤ p<0.001, ¤¤¤¤ p<0.0001 : compared to the first bin or session, or first bin being significantly different from the others

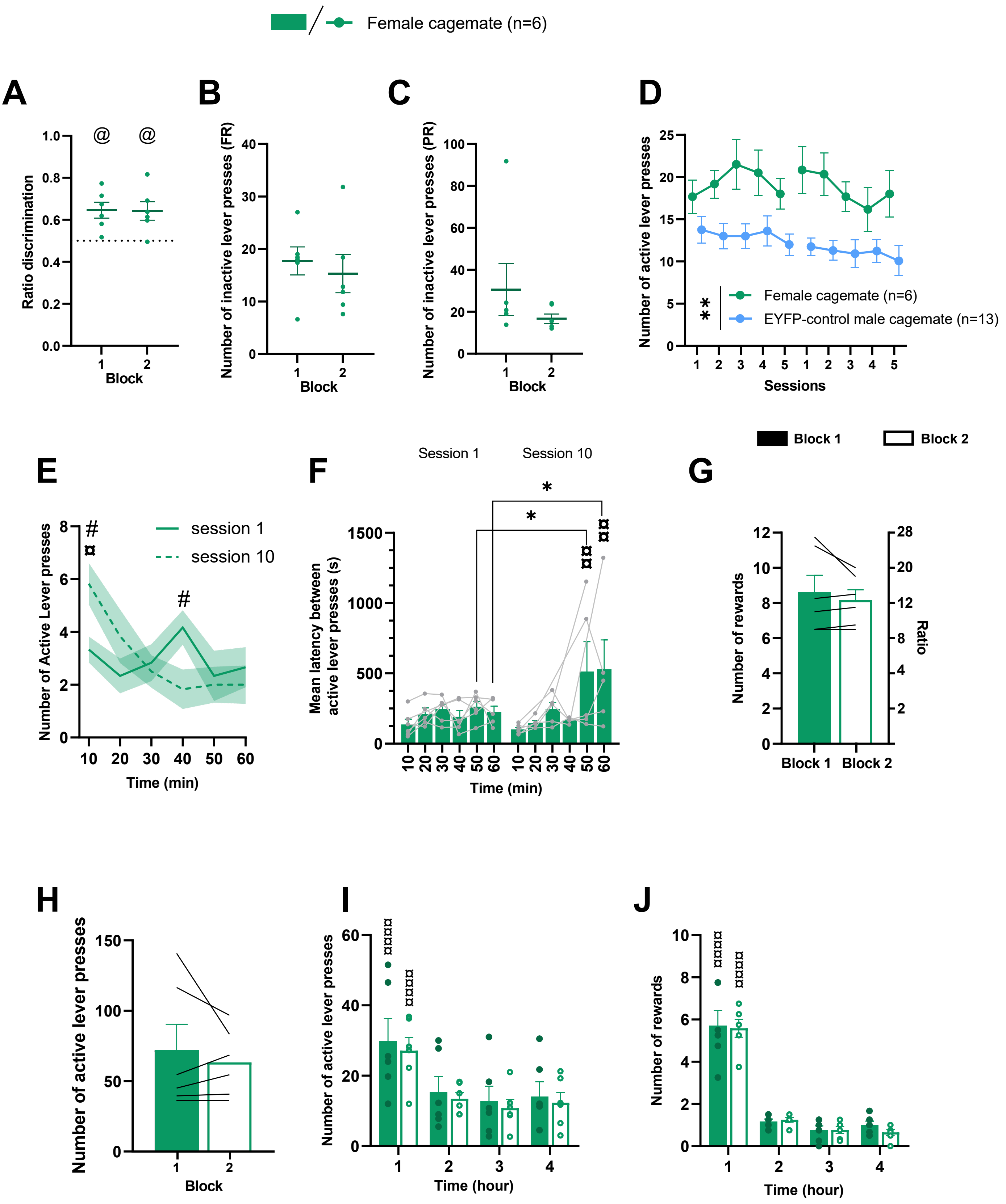

Female cagemate (n=6)
A
Ratio discrimination
Block
@
B
Number of inactive lever presses (FR)
Block
C
Number of inactive lever presses (PR)
Block
D
Number of active lever presses
Sessions
Female cagemate (n=6)
EYFP-control male cagemate (n=13)
**
E
Number of Active Lever presses
Time (min)
session 1
session 10
#
¤
F
Session 1
Session 10
Mean latency between active lever presses (s)
Time (min)
*
¤¤
G
Block 1
Block 2
Number of rewards
Ratio
H
Number of active lever presses
Block
I
Number of active lever presses
Time (hour)
¤¤¤¤
J
Number of rewards
Time (hour)
¤¤¤¤

**Fig. S2 Female rats do not present familiarity-driven reduction in volitional social interactions and motivation**

**(A)** Discrimination ratio in FR1 in female rats (n=6, green) **(B)** Number of inactive lever presses in FR1 (all>0.05) **(C)** Number of inactive lever presses in PR (all>0.05) **(D)** Number of active lever presses per session performed by the female working to interact with cagemate and EYFP-control cagemate males (n=13, light blue) in FR1 (sex : $F(1,17)=11.960$, $p=0.0003$, block: $F(1,150)=8.629$, $p=0.0038$). **(E)** Number of active lever presses per 10 min within session 1 and 10 (bin x session: $F(5,47)=2.877$, $p=0.0223$). **(F)** Mean latency between each lever press per 10 min within session 1 and 10 (bin: $F(5,47)=3.383$, $p=0.0107$). **(G)** Number of social rewards obtained in 4-session blocks in female group in PR (all >0.05). **(H)** Number of active lever presses during block 1 and 2 in PR (all >0.05). **(I) )** Number of active lever presses per 1h in PR (bin : $F(3,175)=3.304$, $p=0.0216$). **(J)** Number of social rewards per 1h in PR (bin : $F(3,164)=29.862$, $p<0.0001$).

@ p<0.0001: significantly different from hazard (50%)

* p<0.05, ** p <0.01, *** p<0.001, **** p<0.0001 : between sex groups

### p<0.05, ## p <0.01, ### p<0.0001 : within sessions or blocks

¤ p<0.05, ¤¤ p <0.01, ¤¤¤ p<0.001, ¤¤¤¤ p<0.0001 : compared to the first bin or session, or first bin being significantly different from the others

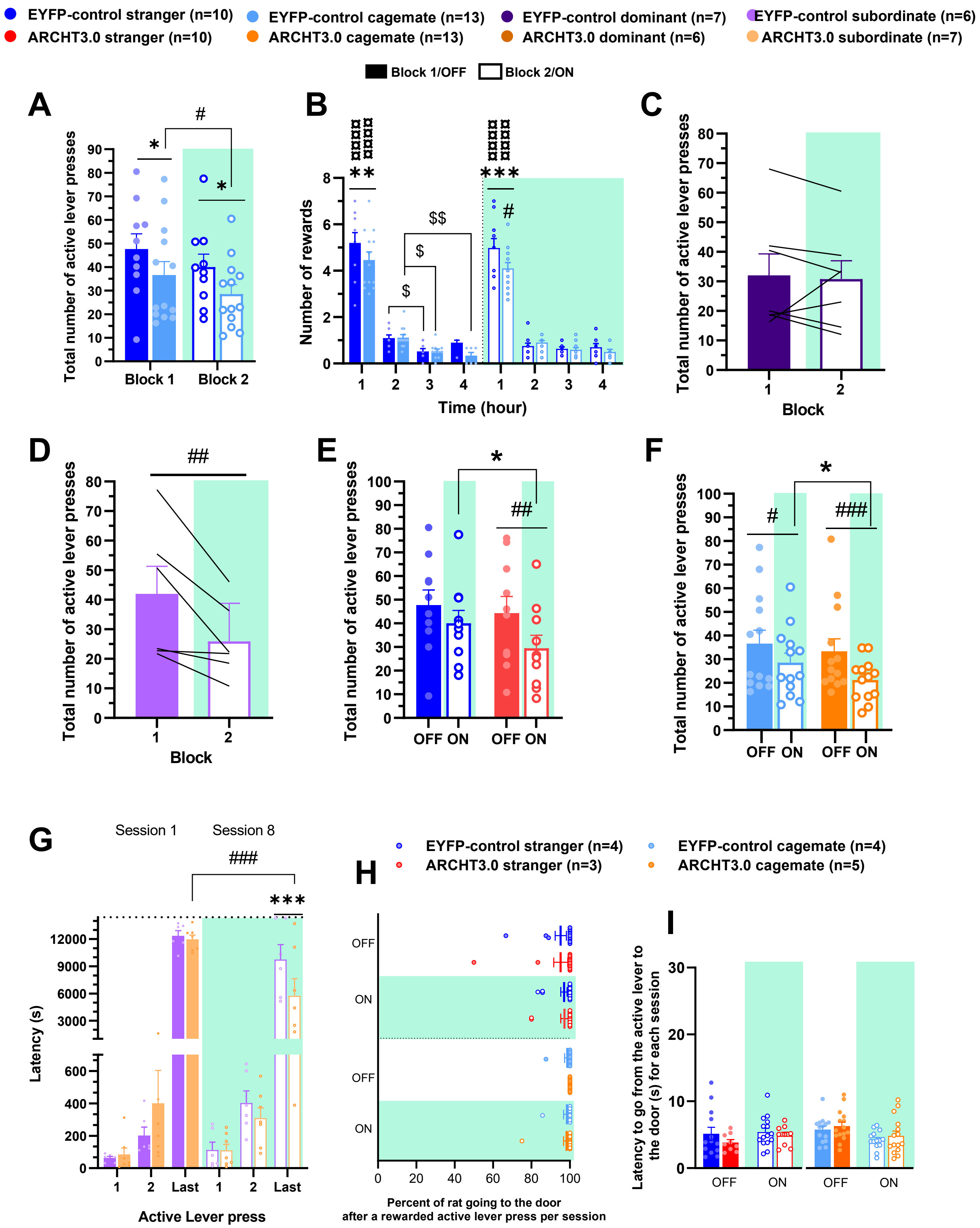
EYFP-control stranger (n=10)
ARCHT3.0 stranger (n=10)
EYFP-control cagemate (n=13)
ARCHT3.0 cagemate (n=13)
EYFP-control dominant (n=7)
ARCHT3.0 dominant (n=6)
EYFP-control subordinate (n=6)
ARCHT3.0 subordinate (n=7)
Block 1/OFF
Block 2/ON
A
Total number of active lever presses
Block 1
Block 2
B
Number of rewards
Time (hour)
C
Total number of active lever presses
Block
D
Total number of active lever presses
Block
E
Total number of active lever presses
OFF ON OFF ON
F
Total number of active lever presses
OFF ON OFF ON
G
Session 1
Session 8
Latency (s)
1
2
Last
1
2
Last
Active Lever press
H
EYFP-control stranger (n=4)
ARCHT3.0 stranger (n=3)
EYFP-control cagemate (n=4)
ARCHT3.0 cagemate (n=5)
OFF
ON
OFF
ON
Percent of rat going to the door
after a rewarded active lever press per session
I
Latency to go from the active lever to
the door (s) for each session
OFF
ON
OFF
ON

**Fig. S3 Familiarity, social hierarchy and role of the STN in males in PR schedule**

**(A)** Total number of active lever presses in EYFP-control groups working to interact with stranger (n=10, dark blue) or with cagemate (n=13, light blue) in PR (block 1: plain bar, block 2: empty bar)(block : $F(1,177)=4.4840$, $p=0.291$, peer identity : $F(1,177)=9.608$, $p=0.0022$). **(B)** Number of social rewards per hour in EYFP-control stranger and cagemate groups (bin x condition : $F(2,592)=7.8418$, $p<0.0001$). **(C)** Total number of active lever presses in EYFP-control dominant (n=7, violet) (all $p>0.05$). **(D)** Total number of active lever presses in EYFP-control subordinate (n=6, pink) (block: $F(1,142)=5.530$, $p=0.0235$). **(E)** Total number of active lever presses in groups working to interact with stranger (ARCHT3.0 stranger (n=10, red) in PR (laser : $F(1,153)=8.306$, $p=0.0045$). **(F)** Total number of active lever presses in groups working to interact with cagemate (ARCHT3.0 cagemate (n=13, orange) in PR (laser : $F(1,201)=15.891$, $p<0.0001$, group : $F(1,201)=4.130$, $p=0.0434$). **(G)** Latency to do the first, second and last lever press in the first (plain bar) and last (empy bar) session in subordinate groups (ARCHT3.0 subordinate, n=7, light orange) (laser x group x active lever press: $F(2,299)=3.606$, $p=0.0283$). **(H)** Percentage of rats that went from the active lever to the door after a rewarded press, without (white area) and with laser (green area), in stranger groups (top, all $p > 0.05$) and in cagemate groups (bottom, all $p>0.05$). **(I)** Latency to go from the active lever to the door after a rewarded press in stranger groups (left, all $p>0.05$) and in cagemate groups (right, all $p>0.05$).

* $p<0.05$, ** $p<0.01$, *** $p<0.001$, **** $p<0.0001$ : between groups (EYFP-control vs ARCHT3.0)

### $p<0.05$, ## $p<0.01$, ### $p<0.0001$ : within sessions or blocks

¤ $p<0.05$, ¤¤ $p<0.01$, ¤¤¤ $p<0.001$, ¤¤¤¤ $p<0.0001$ : compared to the first bin or session, or first bin being significantly different from the others

$ $p<0.05$, $$ $p<0.01$: between bins of the same block

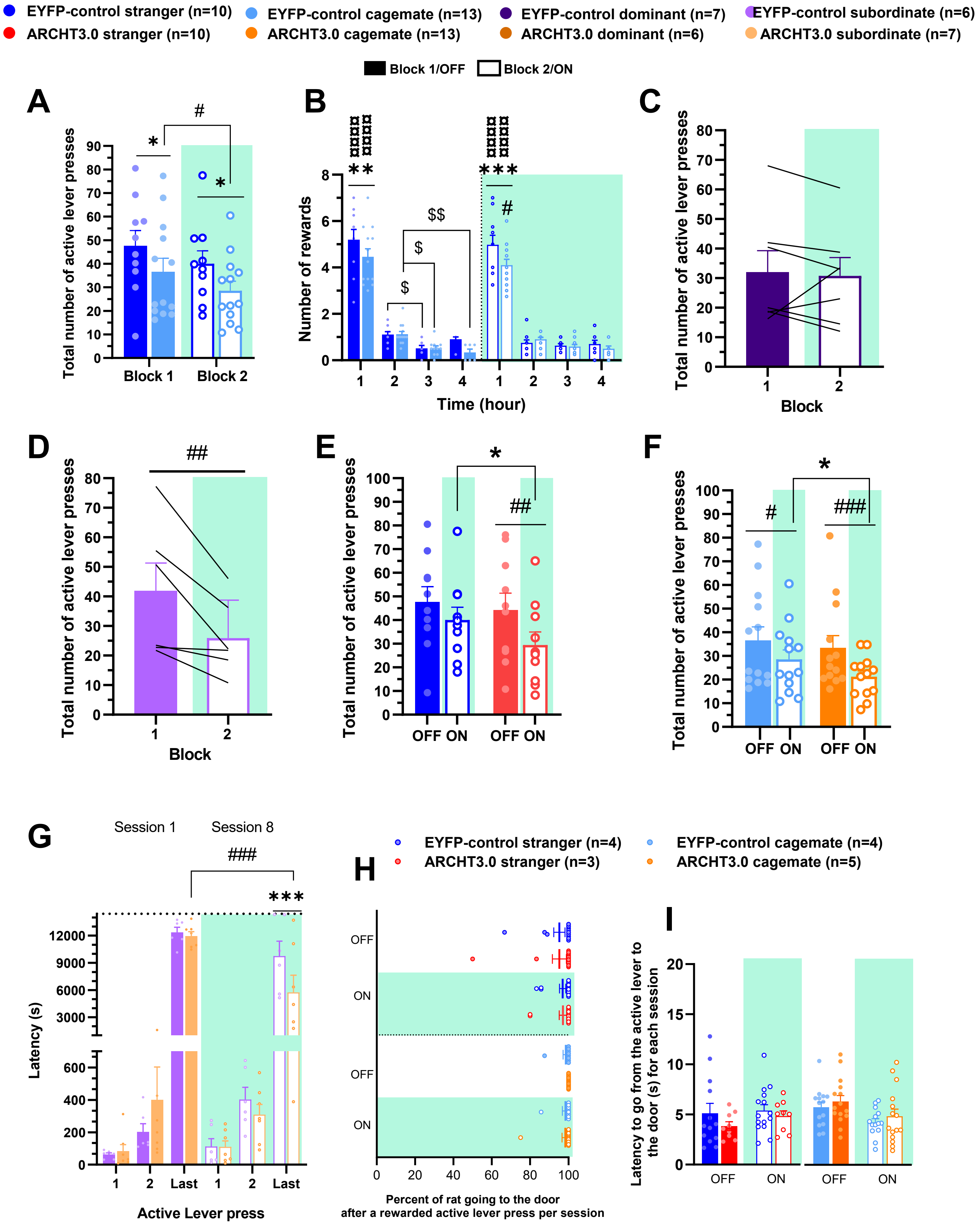
EYFP-control stranger (n=10)
ARCHT3.0 stranger (n=10)
EYFP-control cagemate (n=13)
ARCHT3.0 cagemate (n=13)
EYFP-control dominant (n=7)
ARCHT3.0 dominant (n=6)
EYFP-control subordinate (n=6)
ARCHT3.0 subordinate (n=7)
Block 1/OFF
Block 2/ON
A
Total number of active lever presses
0
10
20
30
40
50
60
70
80
90
*
#
*
Block 1
Block 2
B
Number of rewards
0
2
4
6
8
**
***
#
$
$
$$
1
2
3
4
1
2
3
4
Time (hour)
C
Total number of active lever presses
0
10
20
30
40
50
60
70
80
1
2
Block
D
Total number of active lever presses
0
10
20
30
40
50
60
70
80
##
1
2
Block
E
Total number of active lever presses
0
10
20
30
40
50
60
70
80
90
100
*
##
OFF ON
OFF ON
F
Total number of active lever presses
0
10
20
30
40
50
60
70
80
90
100
*
#
###
OFF ON
OFF ON
G
Session 1
Session 8
###
***
Latency (s)
0
200
400
600
3000
6000
9000
12000
1
2
Last
1
2
Last
Active Lever press
H
EYFP-control stranger (n=4)
ARCHT3.0 stranger (n=3)
EYFP-control cagemate (n=4)
ARCHT3.0 cagemate (n=5)
OFF
ON
OFF
ON
0
20
40
60
80
100
Percent of rat going to the door
after a rewarded active lever press per session
I
Latency to go from the active lever to the door (s) for each session
0
5
10
15
20
OFF
ON
OFF
ON

| Figure | | Effect | F statistic | p-value | Comparison | p-value |
|---|---|---|---|---|---|---|
| 1 | C | Block | **$F_{(1,223)}$=5.451** | **0.0204** | EYFP Stranger vs cagemate:<br>block 1 (session 1- 5)<br>block 2 (session 6-10)<br><br>Block 1 vs 2:<br>EYFP-stranger<br>EYFP-cagemate | <br>0.00181<br><0.0001<br><br><br>0.173<br>0.004 |
| | | peer identity | **$F_{(1,223)}$=26.187** | **<0.0001** | | |
| | | session | $F_{(1,223)}$=0.044 | 0.8336 | | |
| | | block x peer identity | $F_{(1,223)}$=0.252 | 0.6162 | | |
| | D | | | | Session 1 vs 10<br>EYFP-stranger<br>EYFP-cagemate<br><br>Stranger vs Cagemate<br>Session 1<br>Session 10 | <br>0.479<br>0.048<br><br><br>0.352<br>0.0166 |
| | E | 10min_bin | **$F_{(5,143)}$=8.816** | **<0.0001** | EYFP-Cagemate<br>Session 1 :<br>Bin 1 vs 2-5<br>Session 10 :<br>Bin 1 vs 2-6<br><br>Session 1 vs 10 :<br>Bin 4<br>Bin 6 | <br><br>0.0240<br><br><0.039<br><br><br>0.0350<br>0.0440 |
| | | session | **$F_{(1,143)}$=6.640** | **0.0110** | | |
| | | 10min_bin x session | $F_{(5,143)}$=0.782 | 0.5650 | | |
| | F | 10min_bin | $F_{(5,100)}$=1.085 | 0.3735 | EYFP-Cagemate<br>Session 1 :<br>Bin 1 vs 2-3 and 5-6<br>Bin 1 vs 4<br>Session 10 :<br>Bin 1 vs 3-6<br><br>Session 1 vs 10 :<br>Bin 4<br>Bin 6 | <br><br>>0.1252<br>0.0297<br><br><0.0157<br><br><br>0.0160<br><0.0001 |
| | | session | **$F_{(1,103)}$=16.300** | **0.0001** | | |
| | | 10min_bin x session | **$F_{(5,101)}$=4.103** | **0.0020** | | |
| | G | block | **$F_{(1,107)}$=8.034** | **0.0055** | EYFP-subordinate:<br>block 1 vs 2<br>Session 1 vs 10<br><br>EYFP-dominant: block 1 vs 2 | <br>0.0126<br>0.036<br><br>0.514 |
| | | hierarchy | $F_{(1,11)}$=1.091 | 0.3185 | | |
| | | session | $F_{(8,107)}$=0.931 | 0.4946 | | |
| | | block x hierarchy | $F_{(1,107)}$=1.664 | 0.1999 | | |
| | H | 10min_bin | **$F_{(5,66)}$=7.504** | **<0.0001** | EYFP-dominant :<br>Session 1 :<br>Bin 1 vs 2-6<br>Session 10 :<br>Bin 1 vs 2-6<br><br>Session 1 vs 10 :<br>Bin 1 | <br><br>0.7512<br><br><0.0143<br><br><br><0.0143 |
| | | session | $F_{(1,66)}$=0.372 | 0.5438 | | |
| | | 10min_bin x session | $F_{(5,66)}$=2.725 | 0.0267 | | |
| | I | 10min_bin | **$F_{(5,55)}$=7.256** | **<0.0001** | EYFP-subordinate :<br>Session 1 :<br>Bin 1 vs 3-6<br>Session 10 :<br>Bin 1 vs 2-6<br><br>Session 1 vs 10 :<br>Bin 1-3 | <br><br><0.0038<br><br>>0.068<br><br><br><0. 0463 |
| | | session | **$F_{(1,55)}$=18.189** | **<0.0001** | | |
| | | 10min_bin x session | $F_{(5,55)}$=1.024 | 0.4129 | | |
| 2 | A | laser | $F_{(1,193)}$=3.691 | 0.0562 | | |
| | | group | $F_{(1,193)}$=0.086 | 0.7693 | | |
| | | session | $F_{(1,193)}$=0.482 | 0.4885 | | |
| | | laser x group | $F_{(1,193)}$=0.073 | 0.7869 | | |
| | B | laser | $F_{(1,253)}$=3.848 | 0.0509 | EYFP cagemate vs ARCHT3.0 cagemate :<br>block 1 (session 1- 5)<br>block 2 (session 6-10)<br><br>Block 1 vs 2:<br>EYFP-cagemate<br>ARCHT3.0-cagemate | <br><br>0.1970<br>0.0030<br><br><br>0.004<br>0.58 |
| | | group | **$F_{(1,253)}$=9.568** | **0.0022** | | |
| | | session | $F_{(1,253)}$=0.709 | 0.4007 | | |
| | | laser x group | $F_{(1,253)}$=1.777 | 0.1838 | | |
| | C | Bin_10min | **$F_{(5,212)}$=2.544** | **0.029** | EYFP-cagemate vs ARCHT3.0-cagemate | |
| | | group | $F_{(1,48)}$=0.007 | 0.933 | | |

| | | | | | | |
|---|---|---|---|---|---|---|
| | | session | **$F_{(1,217)}$=15.120** | **0.00013** | Session 1 :<br>Bin 1-6<br>Session 10 :<br>Bin 4 and 6<br><br>EYFP-cagemate :<br>Session 1 :<br>Bin 1 vs 4<br>Session 10 :<br>Bin 1 vs 3-6<br>Session 1 vs 10 :<br>Bin 4 and 6<br><br>ARCHT3.0-cagemate<br>Session 1 :<br>Bin 1 vs 6<br>Session 10 :<br>Bin 1 vs 3-4<br>Bin 1 vs 5-6<br>Session 1 vs 10 :<br>Bin 1-6 | <br>0.1524<br><br><0.0177<br><br><br><br>0.0297<br><br><0.0157<br><br><0.0160<br><br><br><br>0.0384<br><br>0.0460<br>>0.3779<br><br>>0.1517 |
| | | Bin_10min x group | $F_{(5,212)}$=0.440 | 0.820 | | |
| | | Bin_10min x session | **$F_{(5,213)}$=3.190** | **0.008** | | |
| | | groupe x session | **$F_{(1,217)}$=10.104** | **0.0017** | | |
| | | Bin_10min x group x session | **$F_{(5,213)}$=3.441** | **0.0052** | | |
| | D | laser | $F_{(1,108)}$=0.232 | 0.6310 | | |
| | | group | $F_{(1,108)}$=0.113 | 0.7370 | | |
| | | session | $F_{(8,108)}$=0.446 | 0.8910 | | |
| | | laser x group | $F_{(1,108)}$=0.113 | 0.7370 | | |
| | E | Bin_10min | F(5, 96) = 1.419 | 0.224 | EYFP-dominant vs ARCHT3.0-dominant<br>Session 1 :<br>Bin 1-6<br>Session 10 :<br>Bin 6<br><br>EYFP-dominant :<br>Session 1 :<br>Bin 1 vs 2-6<br>Session 10 :<br>Bin 1 vs 3-4, 6<br>Session 1 vs 10 :<br>Bin 6<br><br>ARCHT3.0-dominant<br>Session 1 :<br>Bin 1 vs 6<br>Session 10 :<br>Bin 1 vs 4<br>Session 1 vs 10 :<br>Bin 1-6 | <br><br><br>0.3181<br><br>0.0001<br><br><br><br>>0.1806<br><br><0.0267<br><br><0.0001<br><br><br><br>0.6581<br><br>0.0378<br><br>>0.1743 |
| | | group | F(1, 24) = 0.234 | 0.633 | | |
| | | session | **F(1, 98) = 4.979** | **0.028** | | |
| | | Bin_10min × group | F(5, 96) = 0.260 | 0.934 | | |
| | | Bin_10min × session | **F(5, 97) = 2.675** | **0.026** | | |
| | | group × session | F(1, 98) = 3.484 | 0.065 | | |
| | | Bin_10min × group × session | F(5, 97) = 1.826 | 0.115 | | |
| | F | laser | **$F_{(1,107)}$=7.493** | **0.0073** | EYFP subordinate vs ARCHT3.0 subordinate :<br>block 1 (session 1- 5)<br>block 2 (session 6-10)<br><br>Block 1 vs 2<br>EYFP-subordinate<br>ARCHT3.0-subordinate | <br><br>1.0000<br>0.0228<br><br><br>0.0103<br>0.7263 |
| | | group | $F_{(1,11)}$=2.183 | 0.1676 | | |
| | | session | $F_{(8,107)}$=1.317 | 0.2426 | | |
| | | laser x group | **$F_{(1,107)}$=4.648** | **0.0333** | | |
| | G | 10min_bin | $F_{(5,94)}$=1.021 | 0.4097 | EYFP-subordinate vs ARCHT3.0-subordinate<br>Session 1 :<br>Bin 1-6<br>Session 10 :<br>Bin 6<br><br>EYFP-subordinate :<br>Session 1 :<br>Bin 1 vs 2-6<br>Session 10 :<br>Bin 1 vs 4 and 6<br>Session 1 vs 10 :<br>Bin 1 vs 6<br><br>ARCHT3.0-subordinate<br>Session 1 : | <br><br><br>0.3471<br><br><0.0001<br><br><br><br>>0.4651<br><br><0.0366<br><br>0.0001 |
| | | group | $F_{(1,20)}$=0.090 | 0.7666 | | |
| | | session | **$F_{(1,96)}$=8.966** | **0.0035** | | |
| | | 10min_bin x group | $F_{(5,94)}$=0.299 | 0.9120 | | |
| | | 10min_bin x session | $F_{(5,94)}$=0.818 | 0.5396 | | |
| | | group x session | **$F_{(1,96)}$=6.433** | **0.0128** | | |
| | | 10min_bin x group x session | $F_{(5,94)}$=1.331 | 0.2577 | | |

| | | | | | | |
|---|---|---|---|---|---|---|
| | | | | | Bin 1 vs 2-6<br>Session 10 :<br>Bin 1 vs 2-6<br>Session 1 vs 10 :<br>Bin 1-6 | 0.1874<br><br>0.4098<br><br>>0.5314 |
| 3 | A | block | **$F_{(1,177)}$=4.314** | **0.0392** | EYFP-stranger vs EYFP-cagemate:<br>block 1 (session 1- 4)<br>block 2 (session 5-8)<br><br>Block 1 vs 2<br>EYFP-stranger<br>EYFP-cagemate | <br>0.0203<br>0.01<br><br><br>0.116<br>0.048 |
| | | peer identity | **$F_{(1,177)}$=11.855** | **0.0007** | | |
| | | session | $F_{(1,177)}$=0.049 | 0.8248 | | |
| | | block x peer identity | $F_{(1,177)}$=0.002 | 0.9678 | | |
| | B | 1h_bin | **$F_{(3,649)}$=115.967** | **<0.0001** | EYFP-stranger vs EYFP-cagemate<br>Block 1 :<br>Bin 1 and 4<br>Block 2:<br>Bin 1 and 4<br><br>EYFP-stranger<br>Block 1 :<br>Bin 1 vs 2-4<br>Bin 2 vs 3<br>Block 2:<br>Bin 1 vs 2-4<br>Block 1 vs 2 :<br>Bin 1-4<br><br>EYFP-cagemate<br>Block 1 :<br>Bin 1 vs 2-4<br>Bin 2 vs 4<br>Block 2:<br>Bin 1 vs 2-4<br>Bin 2 vs 3-4<br>Block 1 vs 2 :<br>Bin 1-2 | <br><br><0.014<br><br><0.021<br><br><br><br><0.0001<br>0.049<br><br><0.0001<br><br>>0.098<br><br><br><br><0.0001<br>0.0019<br><br><0.0001<br><0.004<br><br>0.046 |
| | | peer identity | $F_{(1,21)}$=3.182 | 0.0888 | | |
| | | block | **$F_{(1,649)}$=11.631** | **0.0007** | | |
| | | session | $F_{(6,649)}$=1.696 | 0.1195 | | |
| | | 1h_bin x peer identity | **$F_{(3,649)}$=6.738** | **0.0002** | | |
| | | 1h_bin x block | $F_{(3,649)}$=1.378 | 0.2485 | | |
| | | peer identity x block | $F_{(1,649)}$=0.398 | 0.5282 | | |
| | | 1h_bin x peer identity x block | $F_{(3,649)}$=0.137 | 0.9378 | | |
| | C | peer identity | $F_{(1,20)}$=1.349 | 0.2588 | EYFP-stranger<br>Block 1 :<br>Reward 1 vs 2-5<br>Block 2:<br>Bin 1 vs 3, 5<br>Block 1 vs Block 2:<br>Reward 3<br><br>EYFP-cagemate<br>Block 1 :<br>Reward 1 vs 4-5<br>Block 2:<br>Bin 1 vs 3-5 | <br><br>>0.2426<br><br><0.0125<br><br>0.0022<br><br><br><br><0.0192<br><br><0.0132 |
| | | block | $F_{(1,816)}$=3.375 | 0.0666 | | |
| | | **Order of the reward** | **$F_{(4,816)}$=12.640** | **<0.0001** | | |
| | | session | $F_{(6,816)}$=0.171 | 0.9847 | | |
| | | peer identity x block | $F_{(1,817)}$=0.015 | 0.9040 | | |
| | | peer identity x Order of the reward | $F_{(4,816)}$=0.247 | 0.9113 | | |
| | | block x Order of the reward | $F_{(4,816)}$=2.184 | 0.0691 | | |
| | | peer identity x block x Order of the reward | $F_{(4,816)}$=0.410 | 0.8013 | | |
| | D | block | $F_{(1,50)}$=0.000 | 1.0000 | | |
| | | session | $F_{(1,50)}$=2.521 | 0.1190 | | |
| | | block x session | $F_{(1,50)}$=0.043 | 0.8370 | | |
| | E | **block** | **$F_{(1,42)}$=5.530** | **0.0235** | EYFP-subordinate<br>Block 1 vs Block 2 | <br>0.004 |
| | | session | $F_{(1,42)}$=0.421 | 0.5199 | | |
| | | block x session | $F_{(1,42)}$=0.044 | 0.8344 | | |
| | F | 1h_bin | **$F_{(3,364)}$=61.028** | **<0.0001** | EYFP-dominant<br>Block 1 :<br>Bin 1 vs 2-4<br>Bin 2 vs 4<br>Block 2:<br>Bin 1 vs 2-4<br>Bin 2 vs 4<br><br>EYFP-subordinate<br>Block 1 :<br>Bin 1 vs 2-4<br>Bin 2 vs 4<br>Block 2:<br>Bin 1 vs 2-4<br>Block 1 vs 2 : | <br><br><0.0007<br>0.0382<br><br><0.0003<br>0.0330<br><br><br><br><0.0002<br>0.0146<br><br><0.0119<br> |
| | | hierarchy | $F_{(1,11)}$=0.060 | 0.8103 | | |
| | | block | **$F_{(1,364)}$=6.350** | **0.0122** | | |
| | | session | $F_{(6,364)}$=1.050 | 0.3925 | | |
| | | 1h_bin x hierarchy | $F_{(3,364)}$=0.169 | 0.9171 | | |
| | | 1h_bin x block | $F_{(3,364)}$=0.660 | 0.5769 | | |
| | | hierarchy x block | **$F_{(1,364)}$=6.698** | **0.0100** | | |
| | | 1h_bin x hierarchy x block | $F_{(3,364)}$=1.127 | 0.3378 | | |

| | | | | | | |
|---|---|---|---|---|---|---|
| | | | | | Bin 1-3 | <0.0317 |
| | G | hierarchy | $F_{(1,10)}=0.022$ | 0.8850 | EYFP-dominant vs EYFP-subordinate<br>Block 2 Reward 5<br><br>EYFP-dominant<br>Block 1 :<br>Reward 1 vs 3-4<br>Block 2:<br>Bin 1 vs 2-5<br><br>EYFP-subordinate<br>Block 1 :<br>Reward 1 vs 2-5<br>Block 2:<br>Bin 1 vs 3-5<br>Block 1 vs 2<br>Reward 3 | <br><br>0.0336<br><br><br><br><0.0347<br><br>>0.0541<br><br><br>>0.1806<br><br><0.0413<br><br><0.0335 |
| | | block | $F_{(1,445)}=3.375$ | 0.0668 | | |
| | | **Order of the reward** | **$F_{(4,445)}=9.209$** | **<0.0001** | | |
| | | session | $F_{(6,445)}=0.738$ | 0.6194 | | |
| | | hierarchy x block | **$F_{(1,445)}=8.242$** | **0.0043** | | |
| | | hierarchy x Order of the reward | $F_{(4,445)}=1.401$ | 0.2327 | | |
| | | block x Order of the reward | $F_{(4,445)}=0.587$ | 0.6725 | | |
| | | hierarchy x block x Order of the reward | $F_{(4,445)}=0.787$ | 0.5341 | | |
| 4 | A | laser | **$F_{(1,153)}=8.306$** | **0.0045** | EYFP-stranger vs ARCHT3.0-stranger :<br>block 1 (session 1- 4)<br>block 2 (session 5-8)<br><br>Block 1 vs 2<br>EYFP-stranger<br>ARCHT3.0-stranger | <br>0.39<br>0.0292<br><br><br>0.116<br>0.0005 |
| | | group | $F_{(1,153)}=0.633$ | 0.4274 | | |
| | | session | $F_{(1,153)}=0.008$ | 0.9284 | | |
| | | laser x group | $F_{(1,153)}=0.872$ | 0.3520 | | |
| | B | 1h_bin | **$F_{(3,552)}=106.151$** | **<0.0001** | EYFP-stranger vs ARCHT3.0-stranger :<br>Block 1<br>Bin 1-4<br>Block 2<br>Bin 1<br><br>EYFP-stranger<br>Block 1 :<br>Bin 1 vs 2-4<br>Bin 2 vs 4<br>Block 2:<br>Bin 1 vs 2-4<br><br>ARCHT3.0-stranger Block 1 :<br>Bin 1 vs 2-4<br>Block 2:<br>Bin 1 vs 2-4<br>Block 1 vs 2 :<br>Bin 1,3,4 | <br>>0.0738<br><br>0.0427<br><br><br><br><0.0001<br>0.0448<br><br><0.0001<br><br><br><br><0.0001<br><br><0.0001<br><br><0.0426 |
| | | laser | **$F_{(1,552)}=12.945$** | **0.0003** | | |
| | | group | $F_{(1,17)}=1.168$ | 0.2941 | | |
| | | session | **$F_{(6,552)}=2.608$** | **0.0168** | | |
| | | 1h_bin x laser | $F_{(3,552)}=0.182$ | 0.9086 | | |
| | | 1h_bin x group | **$F_{(3,552)}=3.181$** | **0.0237** | | |
| | | laser x group | $F_{(1,552)}=3.595$ | 0.0585 | | |
| | | 1h_bin x laser x group | $F_{(3,552)}=0.925$ | 0.4286 | | |
| | C | laser | **$F_{(1,447)}=5.220$** | **0.0228** | EYFP-stranger vs ARCHT3.0-stranger:<br>Block 1 : first, second<br>Last<br>Block 2: first, second, last<br><br>EYFP-stranger:<br>Block 1 vs Block 2<br>First, second, last<br><br>ARCHT3.0-stranger<br>Block 1 vs Block 2<br>First, second<br>Last | <br>>0.9378<br>0.0293<br>0.1562<br><br><br>>0.0866<br><br><br><br>>0.8681<br><0.0001 |
| | | group | $F_{(1,17)}=0.066$ | 0.7995 | | |
| | | Active_lever_press | $F_{(2,447)}=1080.739$ | <0.0001 | | |
| | | session | $F_{(1,447)}=0.088$ | 0.7672 | | |
| | | laser x group | $F_{(1,447)}=2.120$ | 0.1461 | | |
| | | laser x Active_lever_press | **$F_{(2,447)}=9.511$** | **<0.0001** | | |
| | | group x Active_lever_press | $F_{(2,447)}=0.107$ | 0.8984 | | |
| | | laser x group x Active_lever_press | **$F_{(2,447)}=3.037$** | **0.0490** | | |
| | D | laser | **$F_{(1,201)}=15.891$** | **<0.0001** | EYFP-cagemate vs ARCHT3.0-cagemate:<br>block 1 (session 1- 4)<br>block 2 (session 5-8)<br><br>Block 1 vs 2<br>EYFP- cagemate<br>ARCHT3.0- cagemate | <br><br>0.791<br>0.0089<br><br><br>0.048<br><0.0001 |
| | | group | **$F_{(1,201)}=4.130$** | **0.0434** | | |
| | | session | $F_{(1,201)}=0.886$ | 0.3477 | | |
| | | laser x group | $F_{(1,201)}=2.694$ | 0.1023 | | |
| | E | 1h_bin | **$F_{(3,748)}=81.022$** | **<0.0001** | EYFP-cagemate vs ARCHT3.0-cagemate:<br>Block 1<br>Bin 1-4 | <br><br><br>>0.138 |
| | | laser | $F_{(1,748)}=2.503$ | 0.1141 | | |
| | | group | **$F_{(1,748)}=6.570$** | **0.0106** | | |

| | | | | | | |
|---|---|---|---|---|---|---|
| | | session | $F_{(6,748)}$=0.917 | 0.4821 | Block 2<br>Bin 2<br><br>EYFP- cagemate<br>Block 1 :<br>Bin 1 vs 2-4<br>Bin 2 vs 4<br>Block 2:<br>Bin 1 vs 2-4<br>Bin 2 vs 3-4<br>Block 1 vs 2:<br>Bin 1-2<br><br>ARCHT3.0- cagemate Block 1 :<br>Bin 1 vs 2-4<br>Bin 2 vs 3-4<br>Block 2:<br>Bin 1 vs 2-4<br>Bin 2 vs 4<br>Block 1 vs 2 :<br>Bin 2 | <br>0.0083<br><br><br><br><0.0001<br><0.0001<br><br><0.0001<br><0.004<br><br><0.046<br><br><br><0.0002<br><0.0023<br><br><0.0001<br>0.046<br><br>0.0009 |
| | | 1h_bin x laser | $F_{(3,748)}$=1.263 | 0.2860 | | |
| | | 1h_bin x group | $F_{(3,748)}$=1.083 | 0.3555 | | |
| | | laser x group | $F_{(1,748)}$=0.426 | 0.5143 | | |
| | | 1h_bin x laser x group | $F_{(3,748)}$=0.668 | 0.5717 | | |
| | F | laser | $F_{(1,587)}$=0.971 | 0.3248 | EYFP-cagemate vs ARCHT3.0- cagemate:<br>Block 1 : first, second, last<br>Block 2: first, second Last<br><br>EYFP- cagemate:<br>Block 1 vs Block 2<br>First, second, last<br><br>ARCHT3.0- cagemate<br>Block 1 vs Block 2<br>First, second<br>Last | <br>>0.0791<br><br>0.9849<br><0.0001<br><br>>0.2832<br><br><br><br>>0.4085<br><0.0001 |
| | | group | $F_{(1,24)}$=1.369 | 0.2534 | | |
| | | Active_lever_press | **$F_{(2,587)}$=1341.051** | **<0.0001** | | |
| | | session | $F_{(1,587)}$=2.709 | 0.1003 | | |
| | | laser x group | **$F_{(1,587)}$=8.974** | **0.0029** | | |
| | | laser x Active_lever_press | **$F_{(2,587)}$=9.433** | **<0.0001** | | |
| | | group x Active_lever_press | $F_{(2,587)}$=2.162 | 0.1160 | | |
| | | laser x group x Active_lever_press | **$F_{(2,587)}$=7.986** | **0.0004** | | |
| | G | laser | $F_{(1,97)}$=2.552 | 0.1134 | EYFP-dominant vs ARCHT3.0- dominant:<br>block 1 (session 1- 4)<br>block 2 (session 5-8)<br><br>Block 1 vs 2<br>EYFP- dominant<br>ARCHT3.0- dominant | <br>0.701<br>0.0125<br><br><br>1.000<br>0.034 |
| | | **group** | **$F_{(1,97)}$=4.239** | **0.0422** | | |
| | | session | $F_{(1,97)}$=1.663 | 0.2002 | | |
| | | laser x group | $F_{(1,97)}$=2.978 | 0.0876 | | |
| | H | 1h_bin | **$F_{(3,327)}$=379.771** | **<0.0001** | EYFP-dominant vs ARCHT3.0- dominant:<br>Block 1<br>Bin 1-4<br>Block 2<br>Bin 2<br><br>EYFP- dominant<br>Block 1 :<br>Bin 1 vs 2-4<br>Block 2:<br>Bin 1 vs 2-4<br>Bin 3 vs 1-2<br>Block 1 vs Block 2<br>Bin 1-4<br><br>ARCHT3.0- dominant Block 1 :<br>Bin 1 vs 2-4<br>Bin 2 vs 3-4<br>Block 2:<br>Bin 1 vs 2-4<br>Block 1 vs Block 2:<br>Bin 2 | <br><br>>0.205<br><br>0.0372<br><br><br><br><0.0001<br><br><0.0003<br>0.043<br><br>>0.343<br><br><br><0.0001<br><0.040<br><br><0.0001<br><br><0.046 |
| | | group | $F_{(1,11)}$=0.451 | 0.5150 | | |
| | | laser | $F_{(1,328)}$=1.643 | 0.2008 | | |
| | | 1h_bin x group | **$F_{(3,327)}$=3.076** | **0.0278** | | |
| | | 1h_bin x laser | $F_{(3,327)}$=1.365 | 0.2533 | | |
| | | group x laser | $F_{(1,328)}$=0.948 | 0.3309 | | |
| | | 1h_bin x group x laser | $F_{(3,327)}$=0.679 | 0.5656 | | |
| | I | laser | $F_{(1,288)}$=0.721 | 0.3967 | EYFP-dominant vs ARCHT3.0- dominant:<br>Block 1 : first, second, last<br>Block 2: first, second<br>Last<br><br>EYFP- dominant: | <br>>0.0805<br>>0.9408<br>0.0037 |
| | | group | $F_{(1,11)}$=0.107 | 0.7497 | | |
| | | Active_lever_press | $F_{(2,288)}$=534.792 | <0.0001 | | |
| | | session | $F_{(1,288)}$=0.147 | 0.7012 | | |
| | | laser xgroup | **$F_{(1,288)}$=4.113** | **0.0435** | | |

| | | | | | | |
|---|---|---|---|---|---|---|
| | | laser x<br>Active_lever_press | $F_{(2,288)}$=2.453 | 0.0878 | Block 1 vs Block 2<br>First, second, last | >0.6240 |
| | | group x<br>Active_lever_press | $F_{(2,288)}$=0.321 | 0.7261 | ARCHT3.0- dominant | |
| | | laser x group x<br>Active_lever_press | **$F_{(2,288)}$=3.960** | **0.0201** | Block 1 vs Block 2<br>First, second<br>Last | >0.9153<br>0.0008 |
| | J | laser | **$F_{(1,201)}$=15.891** | **<0.0001** | EYFP-subordinate vs ARCHT3.0-<br>subordinate: | |
| | | group | **$F_{(1,201)}$=4.130** | **0.0434** | block 1 (session 1- 4) | 0.8170 |
| | | session | $F_{(1,201)}$=0.886 | 0.3477 | block 2 (session 5-8) | 0.2800 |
| | | laser xgroup | $F_{(1,201)}$=2.694 | 0.1023 | Block 1 vs 2<br>EYFP- subordinate<br>ARCHT3.0- subordinate | 0.004<br>0.0007 |
| | K | Last session (8), one tailed | | | EYFP-subordinate vs ARCHT3.0 subordinate | 0.0406 |
| | L | laser | $F_{(1,299)}$=0.282 | 0.5954 | EYFP- subordinate vs ARCHT3.0-<br>subordinate: | |
| | | group | $F_{(1,299)}$=2.874 | 0.0911 | Block 1 : first, second, last | 0.7708 |
| | | Active_lever_press | $F_{(2,299)}$=848.668 | <0.0001 | Block 2: first, second | 0.9346 |
| | | session | **$F_{(1,299)}$=4.327** | **0.0384** | Last | <0.0001 |
| | | laser x group | **$F_{(1,299)}$=4.506** | **0.0346** | EYFP- subordinate: | |
| | | laser x<br>Active_lever_press | **$F_{(2,299)}$=8.567** | **0.0002** | Block 1 vs Block 2<br>First, second, last | 0.1933 |
| | | group x<br>Active_lever_press | **$F_{(2,299)}$=3.084** | **0.0472** | ARCHT3.0- subordinate | |
| | | laser x group x<br>Active_lever_press | **$F_{(2,299)}$=3.606** | **0.0283** | Block 1 vs Block 2<br>First, second<br>Last | 0.2442<br><0.0001 |

**Table 1: Results summary of statistical analyses of main figures**

| Figure | | Effect | F statistic | p-value | Comparison | p-value |
|---|---|---|---|---|---|---|
| S1 | A | EYFP-control block 1 | **t229 = 13.3** | **<0.0001** | | |
| | | ARCHT3.03.0 block 1 | **t229 = 18.398** | **<0.0001** | | |
| | | EYFP-control block 2 | **t229 = 13.3** | **<0.0001** | | |
| | | ARCHT3.03.0 block 2 | **t229 = 18.398** | **<0.0001** | | |
| | B | block | $F_{(1,193)}=1.312$ | 0.2530 | | |
| | | group | $F_{(1,193)}=0.352$ | 0.5530 | | |
| | | session | $F_{(1,193)}=1.684$ | 0.1960 | | |
| | | block x group | $F_{(1,193)}=0.017$ | 0.8960 | | |
| | C | block | $F_{(1,253)}=0.971$ | 0.3255 | | |
| | | group | $F_{(1,253)}=0.020$ | 0.8872 | | |
| | | session | $F_{(1,253)}=3.411$ | 0.0659 | | |
| | | block x group | $F_{(1,253)}=1.070$ | 0.3019 | | |
| | E | 10min_bin | $F_{(5,50)}=0.756$ | 0.5858 | EYFP-Dominant<br>Session 1<br>Bin 1 vs 2-6<br>Session 10<br>Bin 1 vs 2,4 and 6<br><br>Session 1 vs 10<br>Bin 5 | <br><br>>0.1806<br><br><0.0267<br><br><br><0.0001 |
| | | session | **$F_{(1,51)}=6.023$** | **0.0175** | | |
| | | 10min_bin x session | **$F_{(5,50)}=2.932$** | **0.0212** | | |
| | F | 10min_bin | $F_{(5,39)}=0.359$ | 0.8734 | EYFP-subordinate<br>Session 1<br>Bin 1 vs 2-6<br>Session 10<br>Bin 1 vs 4 and 6<br><br>Session 1 vs 10<br>Bin 5 | <br><br>>0.4651<br><br><0.0366<br><br><br>0.0001 |
| | | session | **$F_{(1,40)}=8.728$** | **0.0052** | | |
| | | 10min_bin x session | $F_{(5,39)}=1.191$ | 0.3309 | | |
| | G | Group- top | $\chi^2(1)= 0.89$, | 0.35 | | |
| | | Group -bottom | $\chi^2(1) < 0.001$ | 1 | | |
| | H | group | $F_{(1,3)}=0.4262$ | 0.5611 | | |
| | | Active_lever_press | $F_{(26,38)}=0.6888$ | 0.8385 | | |
| | | group x Active_lever_press | $F_{(17,38)}=0.8376$ | 6433 | | |
| | I | 10min_bin | **$F_{(5,143)}=8.288$** | **<0.0001** | ARCHT3.0-dominant<br>Session 1<br>Bin 1 vs 2,4-6<br>Session 10<br>Bin 1 vs 3 and 4<br><br>Session 1 vs 10<br>Bin 1-6 | <br><br><0.029<br><br><0.015<br><br><br>>0.07 |
| | | session | $F_{(1,143)}=9.929$ | 0.337 | | |
| | | 10min_bin x session | $F_{(5,143)}=1.331$ | 0.254 | | |
| | J | group | $F_{(1,4)}=0.0121$ | 0.9179 | | |
| | | Active_lever_press | $F_{(18,30)}=1.5004$ | 01576 | | |
| | | group x Active_lever_press | $F_{(12,30)}=0.8376$ | 0.9069 | | |
| S2 | A | block 1 | t34=7.0176 | <0.0001 | | |
| | | block 2 | t24=5.1032 | <0.0001 | | |
| | B | session | $F_{(1,5)}=0.321$ | 0.5950 | | |
| | | bloc | $F_{(1,46)}=1.035$ | 0.3140 | | |
| | | bloc x session | $F_{(1,46)}=0.373$ | 0.5440 | | |
| | C | bloc | $F_{(1,39)}=2.449$ | 0.1257 | | |
| | | session | $F_{(1,39)}=0.220$ | 0.6417 | | |
| | | bloc x session | $F_{(1,39)}=0.493$ | 0.4868 | | |
| | D | Sex | **$F_{(1,17)}=11.960$** | **0.0003** | EYFP-cagemate male vs cagemate female<br>Block 1<br>Block 2<br><br>Block 1 vs Block 2<br>Cagemate Female<br>Cagemate Male EYFP | <br><br><0.0001<br><0.0001<br><br><br>0.497<br>0.004 |
| | | block | **$F_{(1,150)}=8.629$** | **0.0038** | | |
| | | session | $F_{(1,18)}=2.762$ | 0.1140 | | |
| | | block x Sex | $F_{(1,150)}=1.106$ | 0.2946 | | |

| | | | | | | |
|---|---|---|---|---|---|---|
| | E | 10min_bin | **$F_{(5,55)}=2.965$** | **0.0193** | Female cagemate<br>Session 1<br>Bin 1 vs 2-6<br>Session 10<br>Bin 1 vs 3-6<br><br>Session 1 vs 10<br>Bin 1 and 3 | 1.000<br><0.0083<br><0.0243 |
| | | session | $F_{(1,55)}=0.018$ | 0.8930 | | |
| | | 10min_bin x session | **$F_{(5,55)}=2.877$** | **0.0223** | | |
| | F | 10min_bin | **$F_{(5,47)}=3.383$** | **0.0107** | Female cagemate<br>Session 1<br>Bin 1 vs 2-6<br>Session 10<br>Bin 1 vs 5-6<br><br>Session 1 vs 10<br>Bin 5-6 | 1.000<br><0.0068<br><0.0437 |
| | | session | $F_{(1,48)}=1.966$ | 0.1673 | | |
| | | 10min_bin x session | $F_{(5,47)}=1.704$ | 0.1519 | | |
| | G | session | $F_{(1,5)}=2.322$ | 0.1880 | | |
| | | block | $F_{(1,34)}=1.343$ | 0.2550 | | |
| | | block x session | $F_{(1,34)}=0.002$ | 0.9630 | | |
| | H | session | $F_{(1,5)}=0.688$ | 0.4450 | | |
| | | block | $F_{(1,34)}=1.514$ | 0.2270 | | |
| | | block x session | $F_{(1,34)}=0.061$ | 0.8070 | | |
| | I | 1h_bin | **$F_{(3,175)}=3.304$** | **0.0216** | Female cagemate<br>Block 1<br>Bin 1 vs 2-4<br>Block 2<br>Bin 1 vs 2-4<br><br>Block 1 vs 2<br>Bin 1-4 | <0.0001<br><0.0001<br>>0.3503 |
| | | block | $F_{(1,175)}=1.878$ | 0.1723 | | |
| | | session | $F_{(1,175)}=1.874$ | 0.1728 | | |
| | J | 1h_bin | **$F_{(3,164)}=29.862$** | **<0.0001** | Female cagemate<br>Block 1<br>Bin 1 vs 2-4<br>Block 2<br>Bin 1 vs 2-4<br><br>Block 1 vs 2<br>Bin 1-4 | <0.0001<br><0.0001<br>>0.3426 |
| | | block | $F_{(1,164)}=0.085$ | 0.7710 | | |
| | | session | $F_{(6,164)}=0.921$ | 0.4816 | | |
| | | 1h_bin x block | $F_{(3,164)}=0.344$ | 0.7936 | | |
| S3 | A | block | **$F_{(1,177)}=4.840$** | **0.0291** | EYFP-stranger male vs EYFP-cagemate male<br>Block 1<br>Block 2<br><br>Block 1 vs Block 2<br>EYFP-Stranger<br>EYFP-Cagemate | 0.0483<br>0.0141<br><br>0.136<br>0.015 |
| | | peer identity | **$F_{(1,177)}=9.608$** | **0.0022** | | |
| | | session | $F_{(1,177)}=0.036$ | 0.8496 | | |
| | | block x peer identity | $F_{(1,177)}=0.004$ | 0.9523 | | |
| | B | 1h_bin | **$F_{(3,592)}=756.4993$** | **<0.0001** | EYFP-stranger male vs EYFP-cagemate male<br>Block 1<br>Bin 1<br>Block 2<br>Bin 1<br><br>EYFP-stranger<br>Block 1<br>Bin 1 vs 2-4<br>Bin 2 vs 3<br>Block 2<br>Bin 1 vs 2-4<br>Block 1 vs Block 2<br>Bin 1-6<br><br>EYFP-cagemate<br>Block 1<br>Bin 1 vs 2-4<br>Bin 2 vs 3-4<br>Block 2<br>Bin 1 vs 2-4<br>Block 1 vs Block 2<br>Bin 1 | 0.0030<br>0.0005<br><br><0.0001<br>0.0296<br><0.0001<br>>0.0979<br><br><0.0001<br><0.0117<br><0.0001<br>0.0490 |
| | | peer identity | $F_{(1,21)}=3.3292$ | 0.0821 | | |
| | | laser | $F_{(1,592)}=3.6304$ | 0.0572 | | |
| | | session | $F_{(6,592)}=0.6364$ | 0.7012 | | |
| | | 1h_bin x peer identity | **$F_{(3,592)}=7.8418$** | **<0.0001** | | |
| | | 1h_bin x laser | $F_{(3,592)}=1.5046$ | 0.2122 | | |
| | | peer identity x laser | $F_{(1,592)}=0.0679$ | 0.7945 | | |
| | | 1h_bin x peer identity x laser | $F_{(3,592)}=0.2605$ | 0.8539 | | |

| | | | | | | | |
|---|---|---|---|---|---|---|---|
| | C | | block | $F_{(1,50)}=0.000$ | 1.0000 | | |
| | | | session | $F_{(1,50)}=2.521$ | 0.1190 | | |
| | | | block x session | $F_{(1,50)}=0.043$ | 0.8370 | | |
| | D | | block | **$F_{(1,42)}=5.530$** | **0.0235** | EYFP-subordinate<br>Block 1 vs Block 2 | <br>0.004 |
| | | | session | $F_{(1,42)}=0.421$ | 0.5199 | | |
| | | | block x session | $F_{(1,42)}=0.044$ | 0.8344 | | |
| | E | | laser | **$F_{(1,153)}=8.306$** | **0.0045** | EYFP-stranger vs ARCHT3.0-stranger<br>Block 1<br>Block 2<br><br>Block 1 vs Block 2<br>EYFP-Stranger<br>ARCHT3.0-Stranger | <br>0.575<br>0.0495<br><br><br>0.0009<br>0.136 |
| | | | group | $F_{(1,153)}=0.633$ | 0.4274 | | |
| | | | session | $F_{(1,153)}=0.008$ | 0.9284 | | |
| | | | laser x group | $F_{(1,153)}=0.872$ | 0.3520 | | |
| | F | | laser | **$F_{(1,201)}=15.891$** | **<0.0001** | EYFP-cagemate vs ARCHT3.0- cagemate<br>Block 1<br>Block 2<br><br>Block 1 vs Block 2<br>EYFP- cagemate<br>ARCHT3.0- cagemate | <br>0.499<br>0.0359<br><br><br>0.015<br>0.001 |
| | | | group | **$F_{(1,201)}=4.130$** | **0.0434** | | |
| | | | session | $F_{(1,201)}=0.886$ | 0.3477 | | |
| | | | laser x group | $F_{(1,201)}=2.694$ | 0.1023 | | |
| | G | | laser | $F_{(1,299)}=0.282$ | 0.5954 | EYFP-subordinate vs ARCHT3.0- subordinate<br>Session 1<br>First, second, last<br>Session 8<br>First, second<br>Last<br><br>Session 1 vs Session 8<br>EYFP- subordinate<br>First, second, last<br>ARCHT3.0- subordinate<br>Last | <br>>0.6872<br><br>>0.9332<br>0.0004<br><br><br><br><br>0.9764<br><br>0.0001 |
| | | | group | $F_{(1,299)}=2.874$ | 0.0911 | | |
| | | | Active_lever_press | $F_{(2,299)}=848.668$ | <0.0001 | | |
| | | | session | **$F_{(1,299)}=4.327$** | **0.0384** | | |
| | | | laser xgroup | **$F_{(1,299)}=4.506$** | **0.0346** | | |
| | | | laser x Active_lever_press | **$F_{(2,299)}=8.567$** | **0.0002** | | |
| | | | group x Active_lever_press | **$F_{(2,299)}=3.084$** | **0.0472** | | |
| | | | laser x group x Active_lever_press | **$F_{(2,299)}=3.606$** | **0.0283** | | |
| | H | top | laser | $F_{(1,37)}=0.1551$ | 0.6960 | | |
| | | | group | $F_{(1,6)}=0.8109$ | 0.4032 | | |
| | | | session | $F_{(1,37)}=0.7836$ | 0.3818 | | |
| | | | laser x group | $F_{(1,38)}=0.5000$ | 0.4839 | | |
| | | bottom | laser | $F_{(1,53)}=0.5581$ | 0.458 | | |
| | | | groupe | $F_{(1,53)}=2.4180$ | 0.126 | | |
| | | | jour | $F_{(1,53)}=0.7530$ | 0.389 | | |
| | | | laser x group | $F_{(1,53)}=0.5906$ | 0.446 | | |
| | I | Left | Group | $F_{(1,6)}=0.582$ | 0.475 | | |
| | | | Laser | $F_{(1,243)}=0.114$ | 0.736 | | |
| | | | Press_Number | $F_{(7,242)}=0.583$ | 0.769 | | |
| | | | Jour | $F_{(6,243)}=2.082$ | 0.056 | | |
| | | | Group x Laser | $F_{(1,244)}=1.200$ | 0.274 | | |
| | | | Group x Press_Number | $F_{(7,242)}=1.157$ | 0.328 | | |
| | | | Laser x Press_Number | $F_{(7,241)}=0.211$ | 0.983 | | |
| | | | Group x Laser x Press_Number | $F_{(7,241)}=0.335$ | 0.937 | | |
| | | Right | Press_Number | $F_{(7,274)}=1.082$ | 0.375 | | |
| | | | hierarchy | $F_{(1,9)}=0.000$ | 0.989 | | |
| | | | Laser | $F_{(1,278)}=1.573$ | 0.211 | | |
| | | | Jour | $F_{(6,277)}=0.550$ | 0.770 | | |
| | | | Press_Number x hierarchy | $F_{(6,274)}=0.710$ | 0.642 | | |
| | | | Press_Number x Laser | $F_{(7,274)}=0.539$ | 0.804 | | |
| | | | hierarchy x Laser | $F_{(1,279)}=2.143$ | 0.144 | | |
| | | | Press_Number x hierarchy x Laser | $F_{(6,274)}=0.456$ | 0.840 | | |

**Table 2 : Results summary of statistical analyses of supplementals figures**

| | | | | |
|---|---|---|---|---|
| Female: effect of social hierarchy | Number of reward per hierarchy in FR1 | hierarchy | $F_{(1,4)}$=0.077 | 0.7950 |
| | | session | $F_{(4,5)}$=0.958 | 0.3730 |
| | | block | $F_{(1,46)}$=0.530 | 0.4700 |
| | | block x hierarchy | $F_{(1,46)}$=1.683 | 0.2010 |
| | Number of Active lever presses per 10min bin per hierarchy in FR1 | 10min_bin | $F_{(5,44)}$=2.711 | 0.0321 |
| | | hierarchy | $F_{(1,4)}$=0.172 | 0.6993 |
| | | session | $F_{(1,44)}$=0.017 | 0.8978 |
| | | 10min_bin x hierarchy | $F_{(5,44)}$=0.267 | 0.9287 |
| | | 10min_bin x session | $F_{(5,44)}$=2.631 | 0.0364 |
| | | hierarchy x session | $F_{(1,44)}$=1.352 | 0.2512 |
| | | 10min_bin x hierarchy x session | $F_{(5,44)}$=0.721 | 0.6111 |
| | Latencty between each lever presses per 10 min bin per hierarchy in FR1 | 10min_bin | **$F_{(5,36)}$=2.643** | **0.0386** |
| | | hierarchy | $F_{(1,3)}$=0.136 | 0.7322 |
| | | session | $F_{(1,37)}$=1.244 | 0.2718 |
| | | 10min_bin x hierarchy | $F_{(5,36)}$=0.214 | 0.9543 |
| | | 10min_bin x session | $F_{(5,36)}$=1.379 | 0.2549 |
| | | hierarchy x session | $F_{(1,37)}$=0.004 | 0.9506 |
| | | 10min_bin x hierarchy x session | $F_{(5,36)}$=0.721 | 0.6118 |
| | Number of reward per hierarchy in PR | block | $F_{(1,36)}$=0.260 | 0.6135 |
| | | session | $F_{(1,36)}$=0.021 | 0.8851 |
| | | hierarchy | $F_{(1,36)}$=1.069 | 0.3081 |
| | | block x session | $F_{(1,36)}$=0.063 | 0.8029 |
| | | block x hierarchy | $F_{(1,36)}$=0.001 | 0.9723 |
| | | session x hierarchy | $F_{(1,36)}$=0.996 | 0.3248 |
| | | block x session x hierarchy | $F_{(1,36)}$=0.051 | 0.8235 |

**Table 3 : Results summary of statistical analyses of hierarchy influence in female cagemate**
